# Emergence of compensatory mutations reveal the importance of electrostatic interactions between HIV-1 integrase and genomic RNA

**DOI:** 10.1101/2021.12.16.472884

**Authors:** Christian Shema Mugisha, Tung Dinh, Abhishek Kumar, Kasyap Tenneti, Jenna E. Eschbach, Keanu Davis, Robert Gifford, Mamuka Kvaratskhelia, Sebla B. Kutluay

**Affiliations:** Department of Molecular Microbiology, Washington University School of Medicine, Saint Louis, MO 63110, USA; Division of Infectious Diseases, University of Colorado School of Medicine, Aurora, CO 80045; MRC-University of Glasgow Centre for Virus Research, 464 Bearsden Rd., Bearsden, Glasgow G61 1QH, UK

## Abstract

HIV-1 integrase (IN) has a non-catalytic function in virion maturation through its binding to the viral RNA genome (gRNA). Allosteric integrase inhibitors (ALLINIs) and class II IN substitutions inhibit IN-gRNA binding and result in non-infectious viruses marked by mislocalization of the gRNA within virions. HIV-1 IN utilizes basic residues within its C-terminal domain (CTD) to bind to the gRNA. However, the molecular nature of how these residues mediate gRNA binding and whether other regions of IN are involved remain unknown. To address this, we have isolated compensatory substitutions in the background of a class II IN mutant virus bearing R269A/K273A substitutions within the IN-CTD. We found that the nearby D256N and D270N compensatory substitutions restored the ability of IN to bind gRNA and led to the formation of mature infectious virions. Reinstating the local positive charge of the IN-CTD through individual D256R, D256K, D278R and D279R substitutions was sufficient to restore IN-RNA binding and infectivity for the IN R269A/K273A as well as the IN R262A/R263A class II mutants. Structural modeling suggested that compensatory substitutions in the D256 residue created an additional interaction interface for gRNA binding. Finally, HIV-1 IN R269A/K273A, but not IN R262A/R263A, bearing compensatory mutations was more sensitive to ALLINIs providing key genetic evidence that specific IN residues required for RNA binding also influence ALLINI activity. Taken together, our findings highlight the essential role of CTD in gRNA binding and ALLINI sensitivity, and reveal the importance of pliable electrostatic interactions between the IN-CTD and the gRNA.

**IMPORTANCE:** In addition to its catalytic function, HIV-1 integrase (IN) binds to the viral RNA genome (gRNA) through positively charged residues within its C-terminal domain (CTD) and regulates proper virion maturation. Here we show that compensatory mutations in nearby acidic residues (i.e. D256N and D270N) restore the ability to bind gRNA for IN variants bearing substitutions in these positively charged CTD residues. Similarly, charge reversals through individual D-to-R and D-to-K substitutions at these positions enabled the respective IN mutants to bind gRNA and restore virion infectivity. Further, we show that specific residues within the IN-CTD required for RNA binding also influence sensitivity to allosteric integrase inhibitors, a class of novel IN- targeting compounds that target the non-catalytic function of IN. Taken together, our findings reveal the importance of electrostatic interactions in IN-gRNA binding and provide key evidence for a crucial role of the IN-CTD in allosteric integrase inhibitor mechanism of action.

## INTRODUCTION

A defining feature of retroviruses is the reverse transcription of the viral RNA genome (gRNA) and integration of the resultant linear viral DNA into the host chromosome, which establishes lifelong infection. The latter reaction is mediated by the viral integrase (IN) enzyme, which catalyzes 3’ processing and DNA strand transfer reactions (1). The catalytic activity of HIV-1 IN has been successfully targeted by several integrase strand-transfer inhibitors (INSTIs) (2–7) that have become key components of frontline anti-retroviral therapy regimens due to their high efficacy and tolerance profiles (8–11). In addition, HIV-1 IN has a non-catalytic function in virus replication (12–16). Successful targeting of this second function can complement the existing antiviral regimens and substantially increase the barrier to INSTI resistance.

HIV-1 IN consists of three independently folded protein domains: the N-terminal domain (NTD) bears the conserved His and Cys residues (HHCC motif) that coordinate Zn^2+^ binding for 3-helix bundle formation; the catalytic core domain (CCD) adopts an RNase H fold and harbors the enzyme active site composed of an invariant DDE motif, and C-terminal domain (CTD) which adopts an SH3 fold (17, 18). Integration is facilitated by a cellular co-factor, lens epithelium- derived growth factor (LEDGF/p75), which binds tightly to a site within the CCD dimer interface (19, 20) and guides the preintegration complex to actively transcribed regions of the host chromosome (19–24). A group of pleotropic IN substitutions distributed throughout IN, collectively known as class II mutations, disrupt viral assembly (13, 16, 25–36), morphogenesis (12, 16, 27, 32–34, 37, 38) and reverse transcription in target cells (12-14, 16, 29, 31, 32, 34, 36-54) often without obstructing the catalytic activity of IN *in vitro* (13, 27, 28, 31, 40, 41, 44, 46, 55- 57). A hallmark of class II IN mutant viruses is the mislocalization of the viral ribonucleoprotein complexes (vRNP) outside of the viral capsid (CA) lattice, a deformation which is often referred to as eccentric morphology (12, 15, 16, 27, 32–34, 37, 38, 58–60). Although originally designed to inhibit integration through preventing the binding of IN to LEDGF/p75 (58, 61–65), allosteric integrase inhibitors (ALLINIs) potently inhibit proper virion maturation (12, 37, 61) and lead to the formation of virions that display a similar eccentric morphology observed with class II IN mutations (12).

We have recently shown that IN binds to the gRNA in mature virions to mediate proper encapsidation of the viral ribonucleoproteins (vRNPs) inside the mature CA lattice (38). IN binds with high affinity to the TAR hairpin present within the 5’ and 3’UTRs, though multiple other distinct locations on the gRNA without apparent secondary structure are also bound (38). A parameter that appears to be critical for gRNA binding is functional tetramerization of IN. Numerous class II IN mutations scattered throughout IN block IN-gRNA binding indirectly by disrupting functional oligomerization of IN (15). Similar to these class II IN mutations, ALLINIs are thought to interfere with IN-gRNA binding indirectly by inducing aberrant IN multimerization (12, 38). ALLINIs bind to a pocket within the CCD of IN (12, 60, 61, 64, 66–68) and induce the formation of open IN polymers by engaging the CTD of a nearby IN dimer (62). In contrast, mutation of basic residues within IN-CTD (i.e. R262, R263, R269 and K273) inhibits IN-gRNA binding without altering IN oligomerization in virions and *in vitro* (15, 38), suggesting that these residues directly mediate IN binding to the gRNA.

While the involvement of IN-CTD in gRNA binding has been established, how the basic residues within IN-CTD mediate recognition of the gRNA remain unknown. For example, it is possible that the positively charged Lys and Arg residues interact with the negatively charged RNA phosphate backbone in a non-specific or semi-specific manner, depending on the folding and structure of the cognate RNA element, driven by electrostatic interactions (69–71). Alternatively, these residues can mediate specific interactions with gRNA through H-bonding and van der Waals contacts with individual nucleobases (69–71). Furthermore, it is possible that other domains in IN also contribute to gRNA binding. For example, we have previously shown that K34A substitution within the NTD blocks gRNA binding without impacting IN oligomerization (15), suggesting its direct interaction with the gRNA.

To gain insight into the mode of IN-gRNA interactions, the non-infectious HIV-1_NL4-3 IN (R269A/K273A)_ class II IN mutant virus was serially passaged in T-cells until the acquisition of compensatory mutations. Two compensatory substitutions, D256N and D270N, sequentially emerged within the IN-CTD and allowed HIV-1_NL4-3 IN (R269A/K273A)_ to replicate with WT kinetics. Introduction of these substitutions on the HIV-1_NL4-3 IN (R269A/K273A)_ virus backbone substantially increased virion infectivity, restored IN-gRNA binding and proper virion morphogenesis. As the D-to-N mutations resulted in loss of two negative charges, possibly overcoming the loss of two positive charges with the R269A/K273A substitutions, we tested whether restoring the overall local charge of IN-CTD would also restore IN-gRNA binding and virion infectivity. Indeed, the D256R as well as other nearby charge reversal substitutions including D278R and D279R enhanced or restored virion infectivity and RNA binding for the IN R269A/K273A mutant. We further extended these findings to another class II IN mutant, R262A/R263A, which was similarly suppressed by the D256R as well as D256K substitutions. Compensatory mutations did not affect the ability of IN to multimerize *in vitro* or in virions, suggesting that they restored the RNA-binding ability of IN directly. Structural modeling suggested that compensatory mutations created an additional interaction interface for gRNA binding. Finally, the replication ability of the class II IN mutant viruses bearing the compensatory substitutions allowed us to test the contribution of specific CTD residues to ALLINI mechanism of action. Specifically, we found that viruses bearing R269A/K273A, but not the R262A/R263A, substitutions were significantly more sensitive to ALLINIs. Together, our findings highlight the essential role of IN-CTD in gRNA binding, reveal an electrostatic component of IN-gRNA interactions, and provide key genetic evidence that ALLINIs engage specific residues within the IN-CTD.

## RESULTS

### Compensatory IN D256N/D270N substitutions emerge in the background of HIV-1_NL4-3 IN (R269A/K273A)_ class II IN mutant virus

The IN R269A/K273A class II substitutions obstruct IN-gRNA binding directly without interfering with IN multimerization and result in the formation of particles with eccentric morphology (15, 16). To gain insight into the molecular basis of how R269 and K273 residues mediate IN-gRNA binding, HIV-1_NL4-3 IN (R269A/K273A)_ class II IN mutant virus was serially passaged in MT-4 cells until the emergence of compensatory mutations at the end of passage 3, which resulted in virus growth with WT replication kinetics (**Fig. 1A**). Deep sequencing of full-length gRNA isolated from virions across the three passages revealed that the IN R269A and K273A substitutions were retained while the nearby D256N and D270N mutations within IN-CTD were sequentially acquired at the end of passage 1 and passage2, respectively (**Fig. 1B**). Both mutations were fixed by the end of passage 2 and no other non-synonymous mutations were observed in IN or elsewhere on the gRNA.

**Figure 1.**
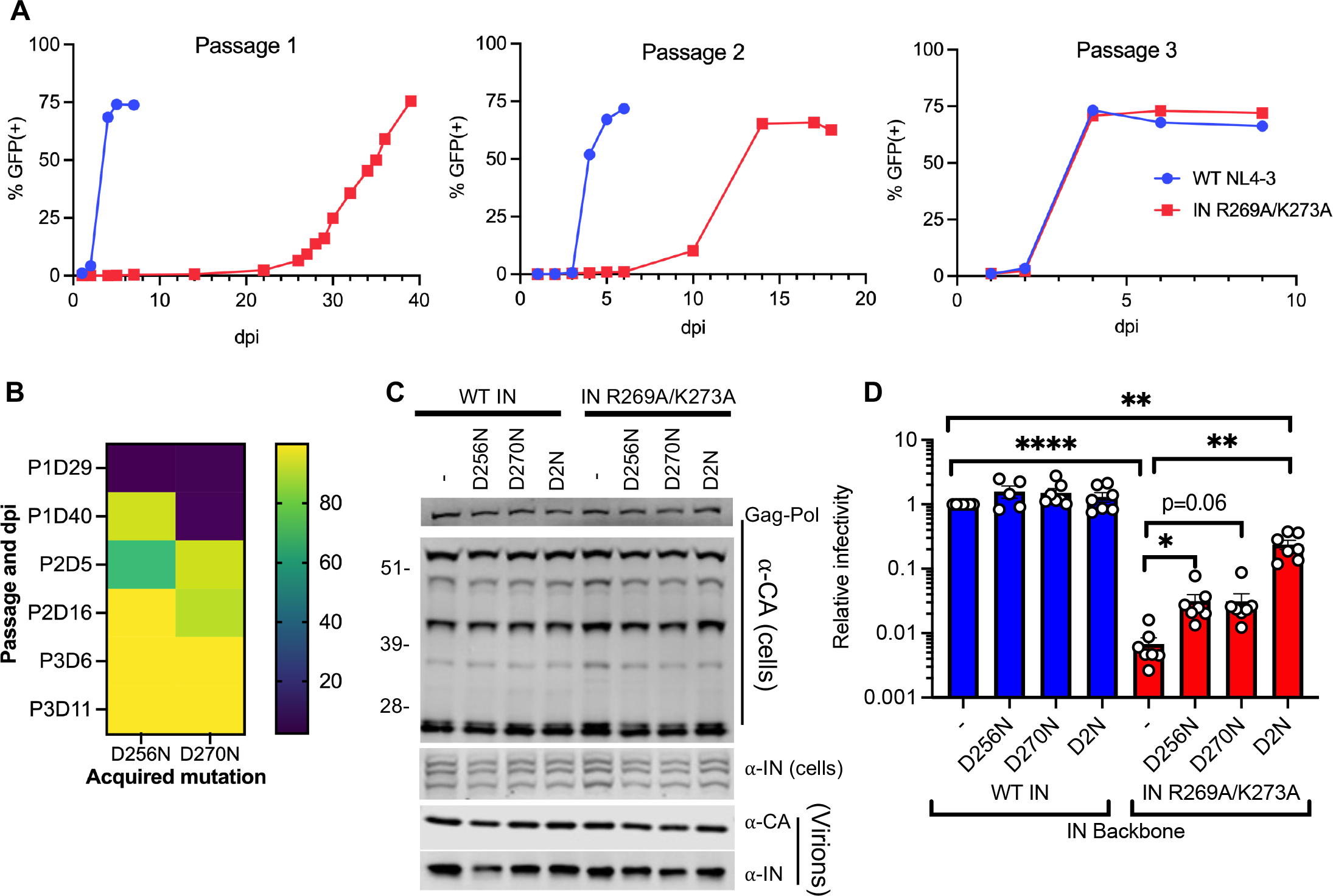
D256N and D270N compensatory substitutions in HIV-1 IN suppress the replication defect of the HIV-1_NL4-3 IN (R269A/K273A)_ class II mutant virus. (A) MT4-LTR-GFP indicator cells were infected with HIV-1_NL4-3_ at an MOI of 2 i.u./cell or an equivalent particle number of the HIV-1_NL4-3 IN (R269A/K273A)_ class II IN mutant virus (based on reverse transcriptase (RT) activity) as detailed in Materials and Methods. HIV-1_NL4-3 IN (R269A/K273A)_ viruses were serially passaged for three rounds until the emergence of compensatory mutations that allowed virus replication with WT kinetics. The graphs represent the percentage of GFP positive cells as assessed by flow cytometry over three passages at the indicated days post-infection (dpi). (B) HIV-1 genomic RNA was isolated from viruses collected from cell culture supernatants over the three passages (i.e. P1, P2, P3) and at the indicated days post-infection (i.e. D29, D40, etc.) as described in Materials and Methods. Heatmap shows the percentage of IN D256N and D270N substitutions at the indicated passages and days post-infection (dpi) as assessed by whole-genome deep sequencing. (C) HEK293T cells were transfected with full-length pNL4-3 expression plasmids carrying *pol* mutations coding for the IN D256N, D270N and D256N/D270N (D2N) substitutions introduced on the WT IN and IN R269A/K273A backbones. Cell lysates and cell culture supernatants containing virions were collected two days post-transfection and analyzed by immunoblotting for CA and IN. The image is representative of five independent experiments. See Fig. S1 for quantitative analysis of immunoblots. (D) HEK293T cells were transfected as in (C) and cell culture supernatants containing viruses were titered on TZM-bl indicator cells, whereby virus replication was limited to a single cycle by addition of dextran sulfate (50 μg/mL). The titers were normalized relative to particle numbers as assessed by an RT activity assay and are presented relative to WT (set to 1). Also see Fig. S1 for titer and RT activity values prior to normalization. The columns represent the average of 5-6 independent biological replicates and the error bars represent standard error of the mean (SEM) (****p<0.0001, **p<0.01, *p<0.05 by one-way ANOVA multiple comparison test with Dunnett’s correction).

The IN D256N and D270N mutations were introduced into the replication-competent pNL4-3 HIV-1 molecular clone bearing WT IN or IN R269A/K273A. Introduction of D256N and D270N substitutions in IN had no observable effect on Gag (Pr55) and Gag-Pol expression or processing in cells, particle release and virion-associated IN levels (**Fig. 1C, S1A-F**). Introduction of D256N and D270N mutations either individually or together (D2N) into HIV-1_NL4-3_ bearing WT IN did not affect viral titers (**Fig. 1D, S1G**). Remarkably, while the individual D256N and D270N mutations introduced into the HIV-1_NL4-3 IN (R269A/K273A)_ class II IN mutant virus increased virus titers by approximately 4-fold, the D2N substitutions increased virus titers by 30- fold (**Fig. 1D, S1G**). Despite no observable differences in multi-cycle replication kinetics (**Fig. 1A**), HIV-1_NL4-3 IN (R269A/K273A/D2N)_ virus was modestly (∼4-fold) but significantly less infectious than WT in single cycle replication assays (**Fig. 1D, S1G**). Overall, these results demonstrate that the combination of D256N and D270N mutations is sufficient to substantially increase the replication competency of the HIV-1_NL4-3 IN (R269A/K273A)_ class II IN mutant virus.

### IN D256N/D270N substitutions restore IN-gRNA binding for the HIV-1_NL4-3 IN (R269A/K273A)_ virus and lead to the formation of correctly matured virions

We next assessed whether the IN D256N and IN D270N substitutions rendered the HIV-1_NL4-3 IN (R269A/K273A)_ virus replication competent by restoring IN-gRNA binding and proper virion maturation. To this end, IN-gRNA complexes were immunoprecipitated from UV-crosslinked virions and visualized per CLIP protocol as described previously (38, 72, 73). IN-RNA complexes were readily visible for viruses bearing WT IN and the D256N, D270N and D2N substitutions introduced on the WT IN backbone did not impact RNA binding (**Fig. 2A, S2**). Expectedly (15, 38), the IN R269A/K273A class II mutant had substantially lower levels of IN-gRNA complexes isolated from virions (**Fig. 2A, S2**). The D256N substitution modestly enhanced the ability of IN R269A/K273A to bind RNA, whereas the D270N substitution had no observable impact (**Fig. 2A, S2**). In contrast, the D2N substitution substantially enhanced the ability of IN R269A/K273A to bind RNA (**Fig. 2A, S2**). The virion morphology of WT and IN mutant viruses was assessed by transmission electron microscopy (TEM). As expected, more than 80% of WT particles had an electron-dense condensate that represents vRNPs inside the CA lattice, whereas the majority of IN R269A/K273A class II mutant virions (∼68%) had a clear eccentric morphology (**Fig. 2B**). Consistent with effects on virus titers and RNA-binding, the introduction of D2N substitutions largely restored the ability of the HIV-1_NL4-3 IN (R269A/K273A)_ virus to form properly matured virions (**Fig. 2B**). Cumulatively, these data show that D256N and D270N substitutions restore infectivity for the HIV-1_NL4-3 IN (R269A/K273A)_ class II IN mutant virus through reestablishing gRNA binding and thereby enabling proper virion maturation.

**Figure 2.**
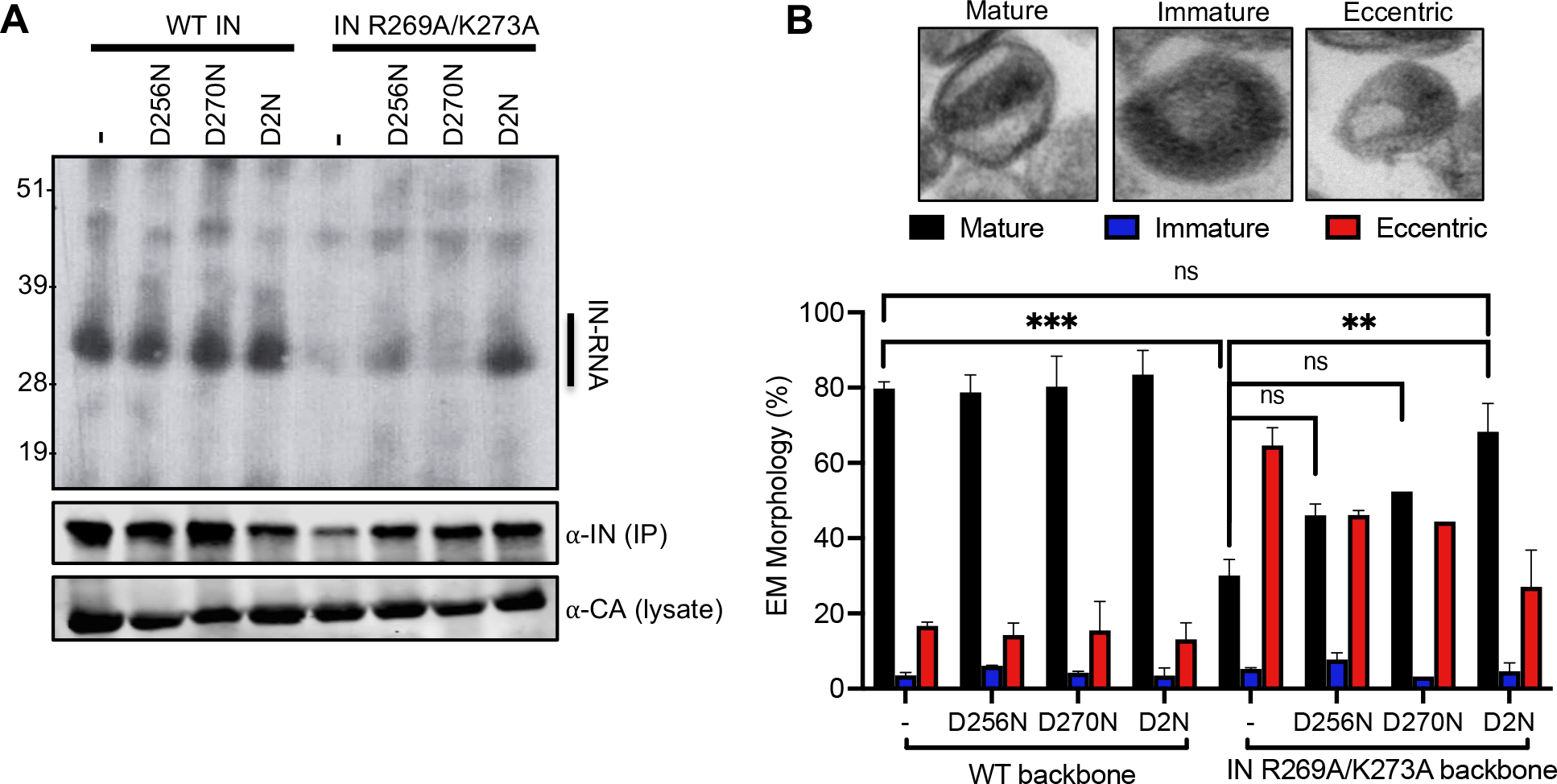
D256N and D270N substitutions restore IN-gRNA binding and accurate virion maturation for the HIV-1_NL4-3 IN (R269A/K273A)_ class II mutant virus. (A) Autoradiogram of IN-RNA adducts immunoprecipitated from virions bearing the indicated substitutions in IN. Immunoblots below show immunoprecipitated (IP) IN or CA protein in lysates prior to immunoprecipitation. Data shown are representative of three independent experiments. See also Fig. S2 for quantitative analysis of autoradiographs. (B) Examination of virion maturation in WT and IN mutant viruses by thin section electron microscopy (TEM). Data show quantification of virion morphologies across 100 particles for each sample and replicate experiment. Data show the average of two independent biological replicates and error bars represent the SEM (***p<0.001, **p<0.01, ns=not significant by one-way ANOVA multiple comparison test with Dunnett’s correction).

### A single charge reversal substitution alone enhances IN-gRNA binding and virion infectivity for the HIV-1_NL4-3 IN (R269A/K273A)_ virus

IN-CTD is decorated with several acidic and basic amino acids, and a long stretch of basic residues spanning 258-273 residues is notable (**Fig. 3A**). In effect, the D2N substitutions restored the overall local charge of this region, suggesting that electrostatic interactions may be a key parameter in gRNA binding. To test this hypothesis, we investigated whether restoring the overall charge of IN-CTD through other mutations would suppress the class II phenotype observed with HIV-1_NL4-3 IN (R269A/K273A)_. We focused our initial analysis on D256 and D270 residues for the following reasons: i) these amino acids were amenable to substitutions during virus passaging experiments; ii) D256N and D270N substitutions in the context of WT HIV-1 yielded infectious virions, suggesting that these compensatory mutations do not significantly contribute to the catalytic activity of IN thus allowing us to specifically probe their roles for IN-gRNA interactions and virion maturation. We introduced D256R, D270R and D256R/D270R (D2R) substitutions into the HIV-1_NL4-3 IN R269A/K273A_ backbone and transfected HEK293T cells with the resulting plasmids. Cell lysates and cell-free virions were then analyzed for Gag and Gag-Pol processing, particle release, and infectivity. Overall, D-to-R substitutions had no major effect on Gag or Gag-Pol expression and processing in cells, or virion release and virion-associated IN levels (**Fig. 3B, S3A-F**).

**Figure 3.**
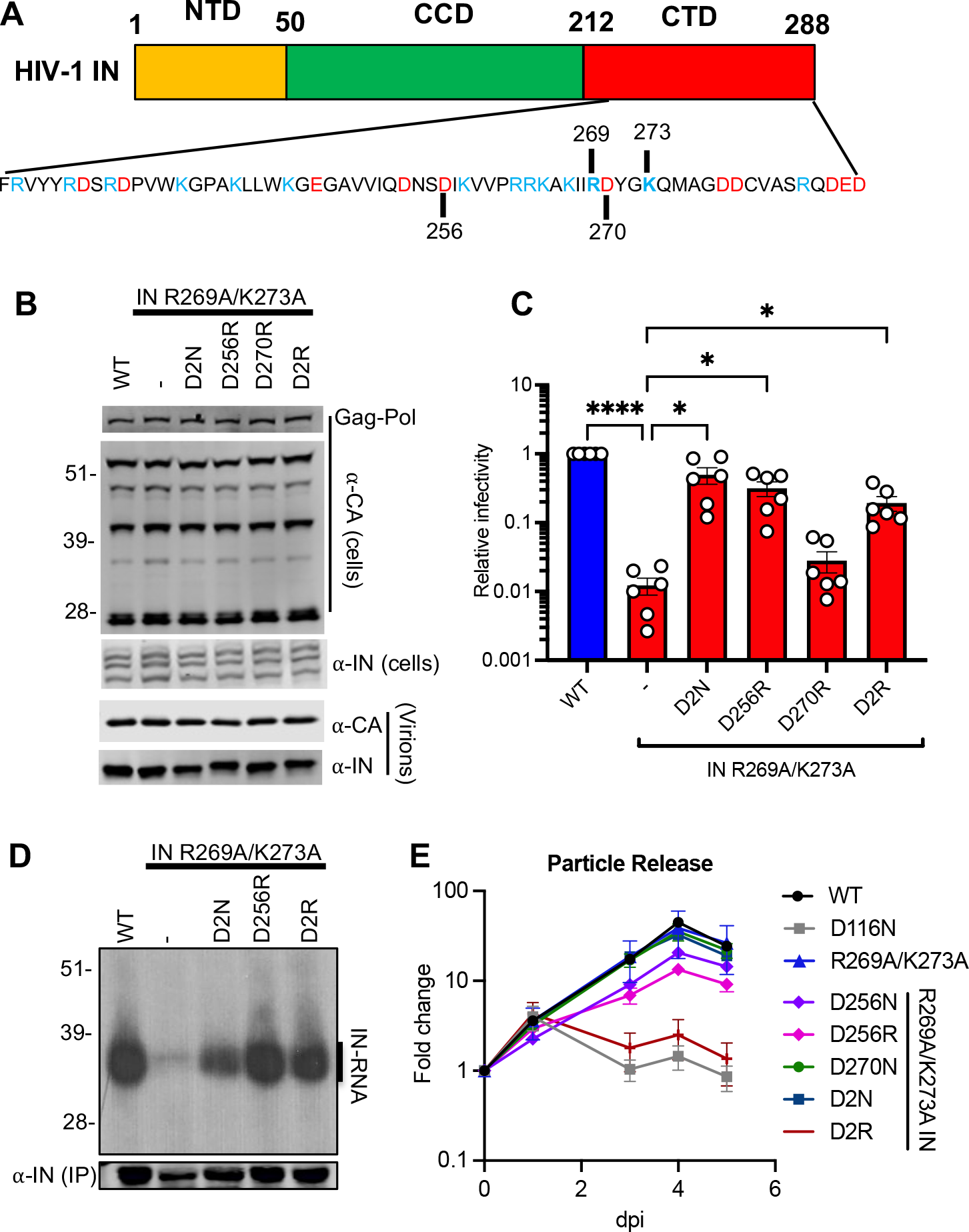
Restoring the local net charge of IN-CTD restores RNA binding and infectivity for the IN R269A/K273A class II mutant. **(A)** Schematic diagram of IN and sequence of CTD residues with basic and acidic amino acids highlighted in blue and red, respectively. **(B, C)** HEK293T cells were transfected with full-length proviral WT HIV-1_NL4-3_ expression plasmid or its derivatives carrying *pol* mutations encoding for D2N, D256R, D270R and D256R/D270R (D2R) IN substitutions on the backbone of HIV-1_NL4-3 IN (R269A/K273A)_. **(B)** Cell lysates and purified virions harvested two days post-transfection were analyzed by immunoblotting for CA and IN. Representative image from five independent experiments is shown. See also Fig. S3A-E for quantitative analysis of immunoblots. **(C)** Cell culture supernatants containing viruses were titered on TZM-bl indicator cells, whereby virus replication was limited to a single cycle by addition of dextran sulfate (50 μg/mL). The titers were normalized relative to particle numbers as assessed by an RT activity assay and are presented relative to WT (set to 1). Also see Fig. S3F, G for titer and RT activity values prior to normalization. The columns represent the average of six independent experiments and the error bars represent SEM (****p<0.0001, *p<0.05 by one-way ANOVA multiple comparison test with Dunnett’s correction). **(D)** Autoradiogram of IN-RNA adducts immunoprecipitated from HIV-1_NL4-3_ virus particles bearing WT IN or the indicated IN mutants. Immunoblot below shows immunoprecipitated (IP) IN protein. Results are representative of three independent replicates. See Fig. S3H for quantitative analysis of autoradiographs. **(E)** MT4 T-cells were infected with HIV-1 _NL4-3 IN (D116N)_ viruses that were trans-complemented with the indicated Vpr-IN mutant proteins as described in Materials and Methods. Virion release was assessed by RT activity assays over the course of 5 days post-infection. Y-axis indicates fold increase in RT activity over day 0. Data are from 3 independent replicates, error bars show the SEM.

Remarkably, the D256R substitution alone increased the titers of HIV-1_NL4-3 IN (R269A/K273A)_ by 26- fold to a level comparable with the D2N substitution (**Fig. 3C, S3G**). In contrast, the D270R substitution did not increase virus titers and the D2R substitution increased the infectivity of HIV- 1_NL4-3 IN (R269A/K273A)_ relatively modestly (∼13-fold) compared to D256R and D2N substitutions (**Fig. 3C, S3G**). Both the D256R and the D2R substitutions restored IN-gRNA binding to WT levels (**Fig. 3D, S3H**), demonstrating that the increase in viral titers with the D256R and D2R substitutions correlates well with enhancement of IN-gRNA binding. To determine whether the D256N/R and D256N/R substitutions affected the catalytic activity of IN, we conducted a transcomplementation assay that relies on incorporation of class II IN mutants into a class I IN mutant virus (HIV-1_NL4-3 IN (D116N)_) bearing a catalytically inactive IN through fusion to the viral Vpr protein (74). In this assay, the RNA-binding defect of a class II IN mutant is complemented *in trans* by the catalytically inactive IN, whereas the class II IN mutant can complement the catalytic activity defect of the class I mutant. We found that none of the IN substitutions impacted the catalytic activity of IN, except for the D2R substitution (**Fig. 3E**), suggesting that the inability of the D270R mutation to restore infectivity to WT levels may be due to its adverse effects on the catalytic activity of IN. Taken together, these findings indicate that restoring the overall charge of IN-CTD through D256R mutation reestablishes virion infectivity and RNA binding for the IN R269A/K273A class II mutant.

### Charge reversal substitutions suppress the replication defect of a separate class II IN mutant virus through restoring gRNA binding

Mutation of other basic residues within the IN-CTD, such as R262A/R263A, also directly inhibit IN-gRNA binding without compromising functional oligomerization of IN (15, 38). If IN-gRNA binding is mediated by electrostatic interactions, we reasoned that substitutions of D256 and D270 residues in a way that restores the local positive charge should restore IN-gRNA binding and infectivity in the background of HIV-1_NL4-3 IN (R262A/R263A)_. D256N, D256K, D256R, D270N and D2N substitutions introduced into HIV-1_NL4-3 IN (R262A/R263A)_ did not have a major impact on Gag and Gag-Pol expression, though a modest increase in distinct Gag processing intermediates was observed (**Fig. 4A, C, S4A-D, S4G-J**). Notwithstanding, neither virion release nor virion-associated IN levels were impacted (**Fig. 4A, C, S4E, S4K, S4L**). IN D256N and D2N substitutions increased virion infectivity by 10- and 5-fold, respectively, whereas the D270N substitution had no impact (**Fig. 4B, S4F**). In contrast, the D256K substitution increased viral titers at a greater degree than D256N and the D256R substitution completely restored virion infectivity (**Fig. 4D, S4M**). In line with the titer data, D256K significantly increased IN-gRNA binding whereas the D256R completely restored it in the context of the HIV-1_NL4-3 IN (R262A/R263A)_ class II mutant virus (**Fig 4E, S4N**). D256N, D256K and D256R IN successfully transcomplemented a class I IN mutant (D116N), suggesting that they did not distort the catalytic activity of IN (**Fig 4F**).

**Figure 4.**
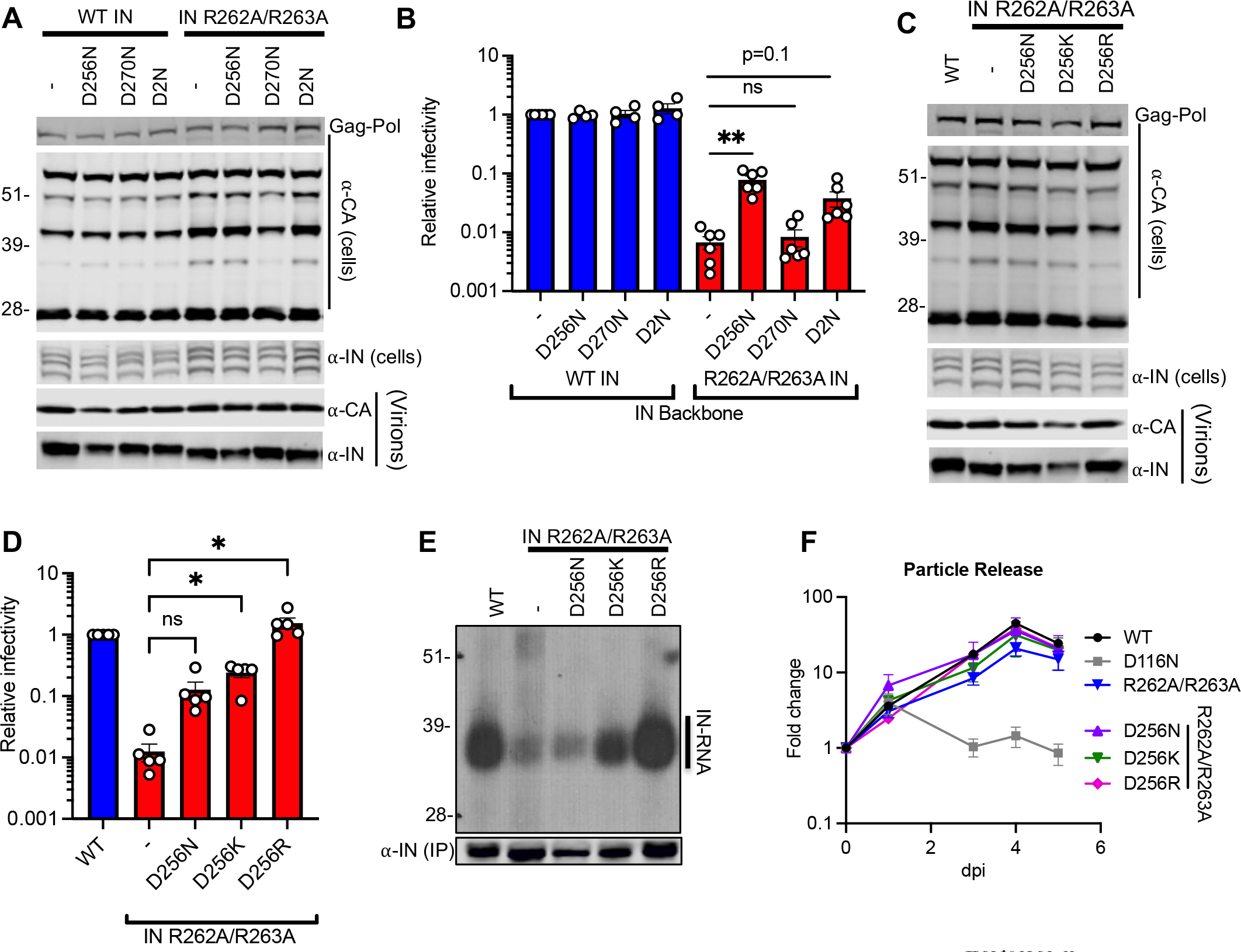
D256R and D256K substitutions restore IN-RNA binding and infectivity for the HIV-1_NL4-3 IN (R262A/R263A)_ class II mutant virus. (A-E) HEK293T cells were transfected with proviral HIV-1_NL4-3_ expression plasmids carrying *pol* mutations for the indicated IN substitutions on WT IN or IN R262A/R263A backbones. (A, C) Cell lysates and purified virions collected two days post-transfection were analyzed by immunoblotting for CA and IN. See also Fig. S4 for quantitative analyses of immunoblots. (B, D) Cell culture supernatants containing viruses were titered on TZM-bl indicator cells, whereby virus replication was limited to a single cycle by addition of dextran sulfate (50 μg/mL). The titers were normalized relative to particle numbers as assessed by an RT activity assay and are presented relative to WT (set to 1). See also Fig. S4F, L, M for titer and RT activity values prior to normalization. The columns represent the average of four-to-six independent experiments, and the error bars represent SEM (**p<0.01, *p<0.05, ns=not significant by one-way ANOVA multiple comparison test with Dunnett’s correction). (E) Autoradiogram of IN-RNA adducts immunoprecipitated from WT or IN mutant HIV-1_NL4-3_ virions. The amount of immunoprecipitated IN was assessed by the immunoblot shown below. Immunoblots and CLIP autoradiographs are representative of three independent replicates. See Fig. S4N for quantitative analysis of autoradiographs. (F) MT4 T-cells were infected with HIV-1 _NL4-3 IN (D116N)_ viruses that were trans-complemented with the indicated Vpr-IN mutant proteins as described in Materials and Methods. Virion release was assessed by RT activity assays over the course of 5 days post-infection. Y-axis indicates fold increase in RT activity over day 0. Data are from 3 independent replicates, error bars show the SEM.

### Charge reversal substitutions at acidic residues other than D256 can suppress the class II phenotype

Given the above findings, we next determined whether charge reversal substitutions at other nearby acidic residues could restore IN-gRNA binding and infectivity for the HIV-1_NL4-3 IN (R269A/K273A)_ virus. Remarkably, the D279R substitution increased virus infectivity by 18-fold and the D278R/D279R substitutions restored infectivity to WT levels (**Fig. 5A**). We also determined whether substitutions of the original D256 and D270 residues into Ile would restore virus titers at levels similar to the D-to-N substitutions. However, we found that neither D256I alone nor the D256I/D270I (D2I) substitutions increased viral titers (**Fig. 5A**). Of note, these substitutions did not impact Gag and Gag-Pol expression, processing, virion release or virion-associated IN levels, though we noted the presence of an aberrantly processed IN with the D2I mutant (**Fig. 5B**). In line with the titer data, D279R substitution increased and the D2R substitution restored IN-RNA binding for the HIV-1_NL4-3 IN (R269A/K273A)_ virus (**Fig. 5C**). Taken together, these results demonstrate that restoring the local positive charge of IN-CTD is sufficient to restore virion infectivity and RNA binding to two distinct class II IN mutants, which suggests the importance of electrostatic interactions in mediating IN-gRNA binding.

**Figure 5.**
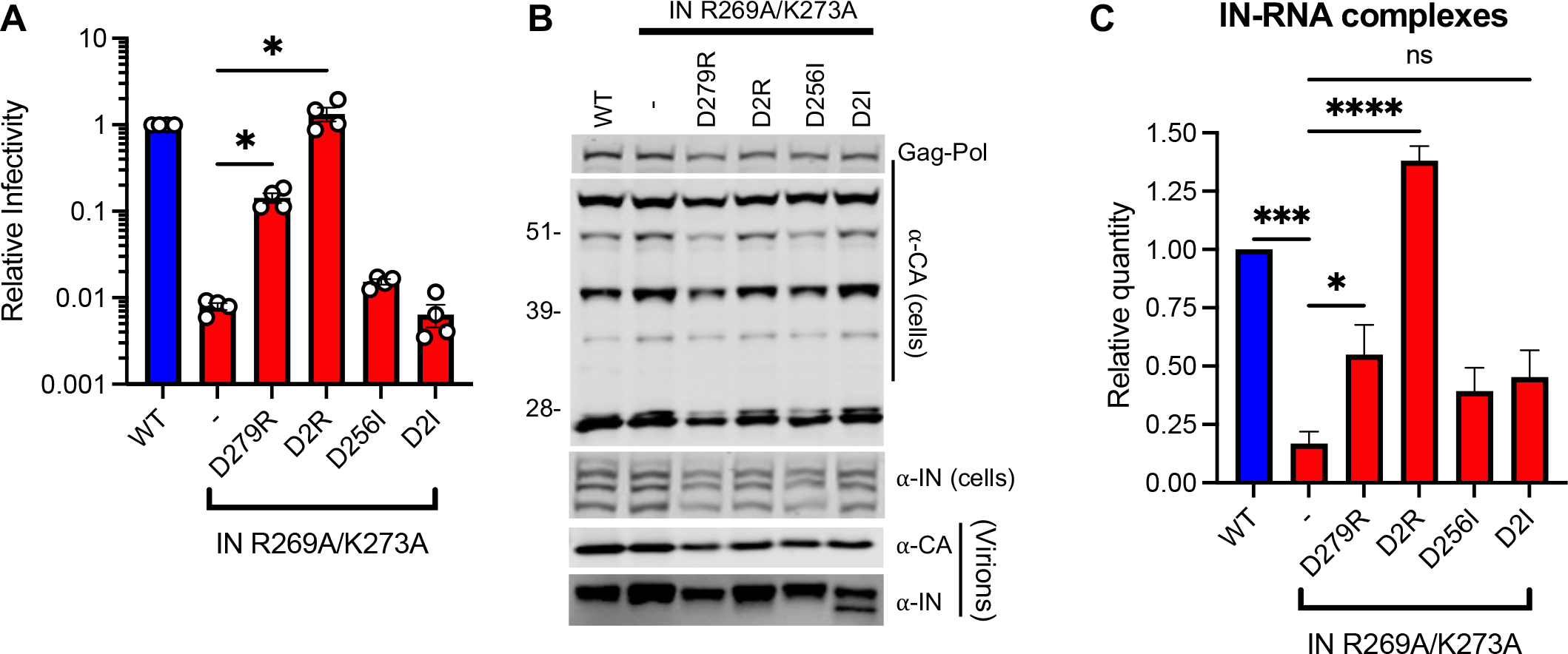
D279R substitution increases and D278R/D279R substitutions restore IN-RNA binding and infectivity for the HIV-1_NL4-3 IN (R269A/K273A)_ virus. HEK293T cells were transfected with full-length pNL4-3 expression plasmids carrying mutations coding for the IN D279R, D278R/D279R (D2R), D256I and D256I/D270I (D2I) substitutions introduced on the IN R269A/K273A backbone. (A) Cell culture supernatants containing viruses were titered on TZM- bl indicator cells, whereby virus replication was limited to a single cycle by addition of dextran sulfate (50 μg/mL). The titers were normalized relative to particle numbers by an RT activity assay and are presented relative to WT (set to 1). The columns represent the average of 4 independent biological replicates and the error bars represent standard error of the mean (SEM) (*p<0.05 by one-way ANOVA multiple comparison test with Dunnett’s correction). (B) Cell lysates and purified virions collected two days post transfection were analyzed by immunoblotting for CA and IN. The image is representative of four independent experiments. (C) IN-RNA adducts immunoprecipitated from WT or IN mutant HIV-1_NL4-3_ virions per CLIP protocol were analyzed by autoradiography. Graph shows the quantification of IN-RNA adducts from three independent biological replicates, error bars show the SEM.

### Compensatory substitutions create an alternative interaction surface for RNA binding

We next used reported X-ray structure (62) and molecular modelling to visualize how class II and compensatory IN mutations affect the electrostatic potential of the CTD surface. R269A and K273A substitutions expectedly resulted in a substantial loss of a basic patch in IN (**Fig. 6A, B**). Both the D256R and D256N substitutions resulted in a more positively charged surface distal from the R269/K273A residues (**Fig. 6C, D**), suggesting that compensatory mutations likely created an additional interaction interface for gRNA binding. A similar outcome was observed when class II R262A/R263A and compensatory D256R changes were introduced in the CTD of IN (**Fig. 6E-G**). While the D256R change is substantially distanced from R262/R263 and R269/K273 residues, we note the following. R262, R263, R269, and K273 are positioned within the same highly flexible C-terminal tail (aa 261-275), whereas D256 belongs to another, shorter (aa 252-257) loop (**Fig. 6H**). The highly pliable nature of the tail and the loop could be crucial for IN to optimally engage with cognate RNA as well as allow for emergence of compensatory mutations at alternative sites positioned in these IN segments.

**Figure 6.**
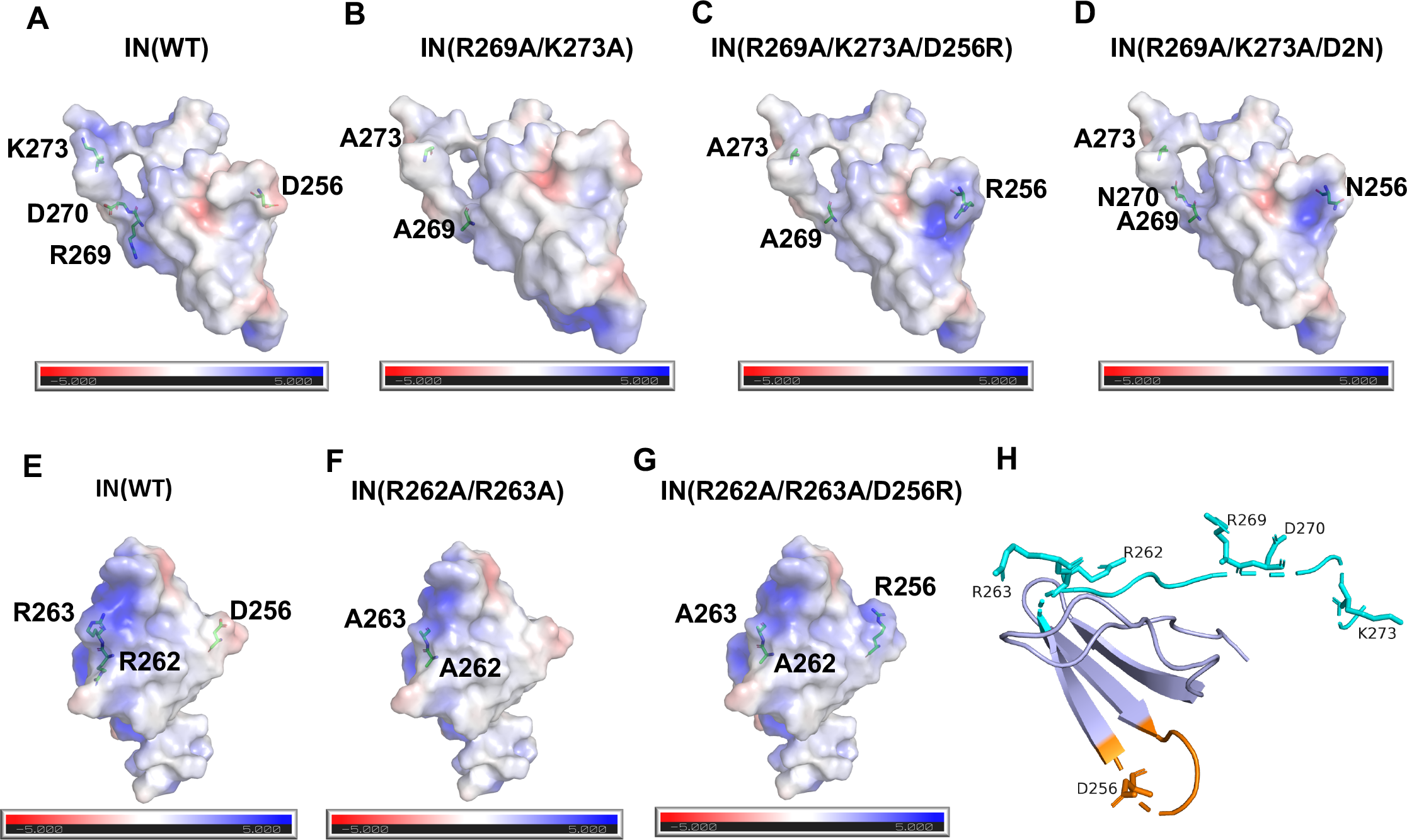
Electrostatic potential maps of HIV-1 IN bearing class II and compensatory mutations. **(A-G)** Electrostatic potential maps of the indicated HIV-1 IN mutants derived from the crystal structure of Gupta et al. (62) is depicted. Results are displayed as an electrostatic potential molecular surface. The low, mid, and high range values are -5, 0, and 5, respectively. (**H**) A cartoon view of the CTD structure (62) is shown with the C-terminal tail (aa 261-275) and the loop (aa 252-257) colored in cyan and orange, respectively. Side chains of indicated residues are shown.

### Compensatory mutations directly restore the ability of IN to bind vRNA without altering functional IN oligomerization

As functional IN oligomerization is a prerequisite for RNA binding (15), we next examined how the compensatory substitutions affected IN oligomerization. For in virion analysis, purified HIV- 1_NL4-3_ virions (WT and bearing the relevant IN substitutions) were treated with ethylene glycol bis (succinimidyl succinate) (EGS) to covalently crosslink IN in situ and virus lysates were analyzed by immunoblotting. As in WT virions, IN species that migrated at molecular weights consistent with those of monomers, dimers, trimers, and tetramers were readily distinguished in HIV-1_NL4-3 IN (R269A/K273A)_ and HIV-1_NL4-3 IN (R262A/R263A)_ viruses with additional compensatory mutations (D256N/D270N, D256R, D256K, and D256R/D270R) but not with the canonical class II IN mutant V165A that is unable to form functional oligomers (**Fig 7A-C** & (15)). Complementary in vitro assessment of purified recombinant IN proteins by size exclusion chromatography (SEC) revealed that D2N and D256R substitutions in the background of IN R269A/K273A or the IN R262A/R263A class II mutants did not impact functional IN tetramerization (**Fig. 7D-H**).

**Figure 7.**
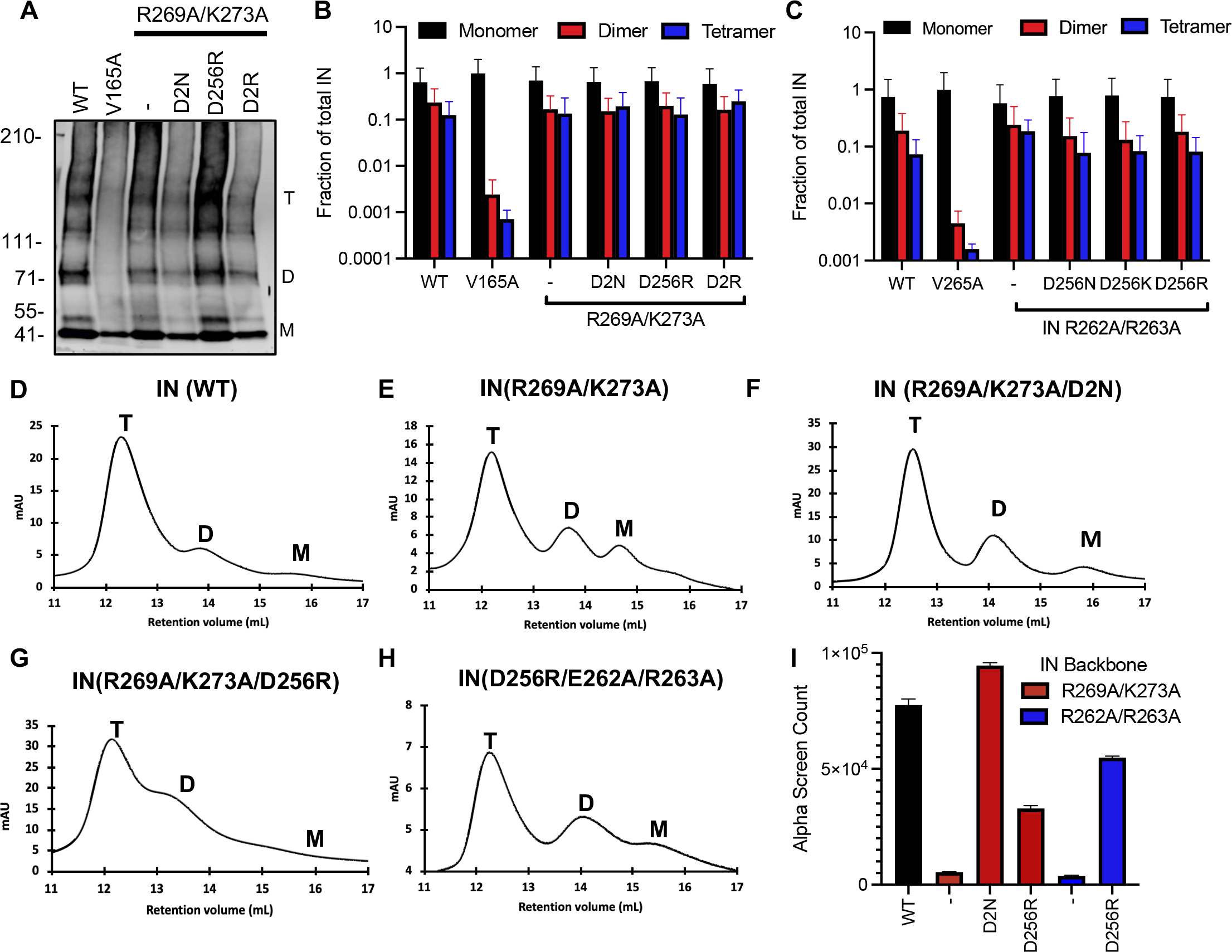
Assessing multimerization properties of IN in mutant viruses. **(A)** Purified WT or IN mutant HIV-1_NL4-3_ virions were treated with 1 mM EGS, and virus lysates analyzed by immunoblotting using antibodies against IN following separation on 3-6% Tris-acetate gels as detailed in Materials and Methods. The position of monomers (M), dimers (D), and tetramers (T) are indicated by arrows in a representative western blot. **(B, C)** Quantification of IN multimerization in virions from experiments conducted as in A. Error bars show the SEM from three independent experiments. **(D-H)** Representative SEC traces for indicated recombinant IN proteins. The X-axis indicates elution volume (mL) and Y-axis indicates the intensity of absorbance (mAU). Tetramers (T), dimers (D), and monomers (M) are indicated. **(I)** Summary of mutant INs bridging TAR RNA compared to WT IN. Alpha screen counts at 320 nM for each protein is shown. The graphs show average values of three independent experiments and the error bars indicate standard deviation.

We have previously shown that recombinant IN binds to TAR RNA with high affinity and provides a nucleation point to bridge and condense RNA (38). We next examined the ability of class II IN mutants bearing compensatory mutations to bind and bridge TAR RNA. Consistent with findings from CLIP, the D2N and D256R substitutions also enhanced or restored the ability of IN R269A/K273 and IN R262A/R263A mutants to bridge between RNA molecules (**Fig. 7I**). Together, these data demonstrate that the compensatory mutations directly restore the ability of IN to bind RNA without altering functional IN oligomerization.

### Sensitivity to ALLINIs is determined by distinct residues within the CTD

ALLINIs potently disrupt proper virion maturation through inducing aberrant IN multimerization and consequently inhibiting IN-gRNA binding (12, 38). Recent structural studies have shown that the quinoline-based ALLINI, GSK-1264, can directly engage residues within the IN-CTD through its tert-butoxy and carboxylic acid moieties and induce open IN polymers (62). These findings are in line with previous biochemical and modeling studies that also showed the involvement of the IN-CTD, in particular residues K264 and K266, in ALLINI-induced aberrant IN multimerization (75, 76). Based on these prior findings, we wanted to test how adjacently positioned IN R262A/R263A and R269A/K273A substitutions and the compensatory mutations affected the antiviral activities of ALLINI.

To this end, we examined effects of representative quinoline-based ALLINIs BI-D and BI-B2, which share many chemical features with GSK-1264, on the viruses bearing the class II and compensatory IN mutations. While the titers of WT viruses only decreased at ALLINI concentrations greater than 1 μM, the titers of HIV-1_NL4-3 IN (R269A/K273A)_ viruses bearing D2N, D256R and D2R substitutions were reduced by 10-fold at ALLINI concentrations as low as 0.1 μM (**Fig. 8A, B**). In contrast, viruses bearing IN R262A/R263A/D256R were less sensitive to low concentrations of ALLINIs (**Fig. 8D, E**), suggesting that the IN R269A and/or K273A, but not the compensatory substitutions, underlie the increased sensitivity to ALLINIs. The increased sensitivity to ALLINIs correlated well with inhibition of RNA binding. While 0.1 μM of BI-D didn’t affect the level of IN-gRNA binding in WT HIV-1, it significantly reduced IN-gRNA binding in HIV-1_NL4-3 IN (R269A/K273A)_ but not HIV-1_NL4-3 IN (R262A/K263A)_ bearing compensatory mutations (**Fig. 8C,F**). Taken together, our findings provide key genetic, biochemical and virological evidence that specific CTD residues required for RNA binding are also crucial for the ALLINI mechanism of action.

**Figure 8.**
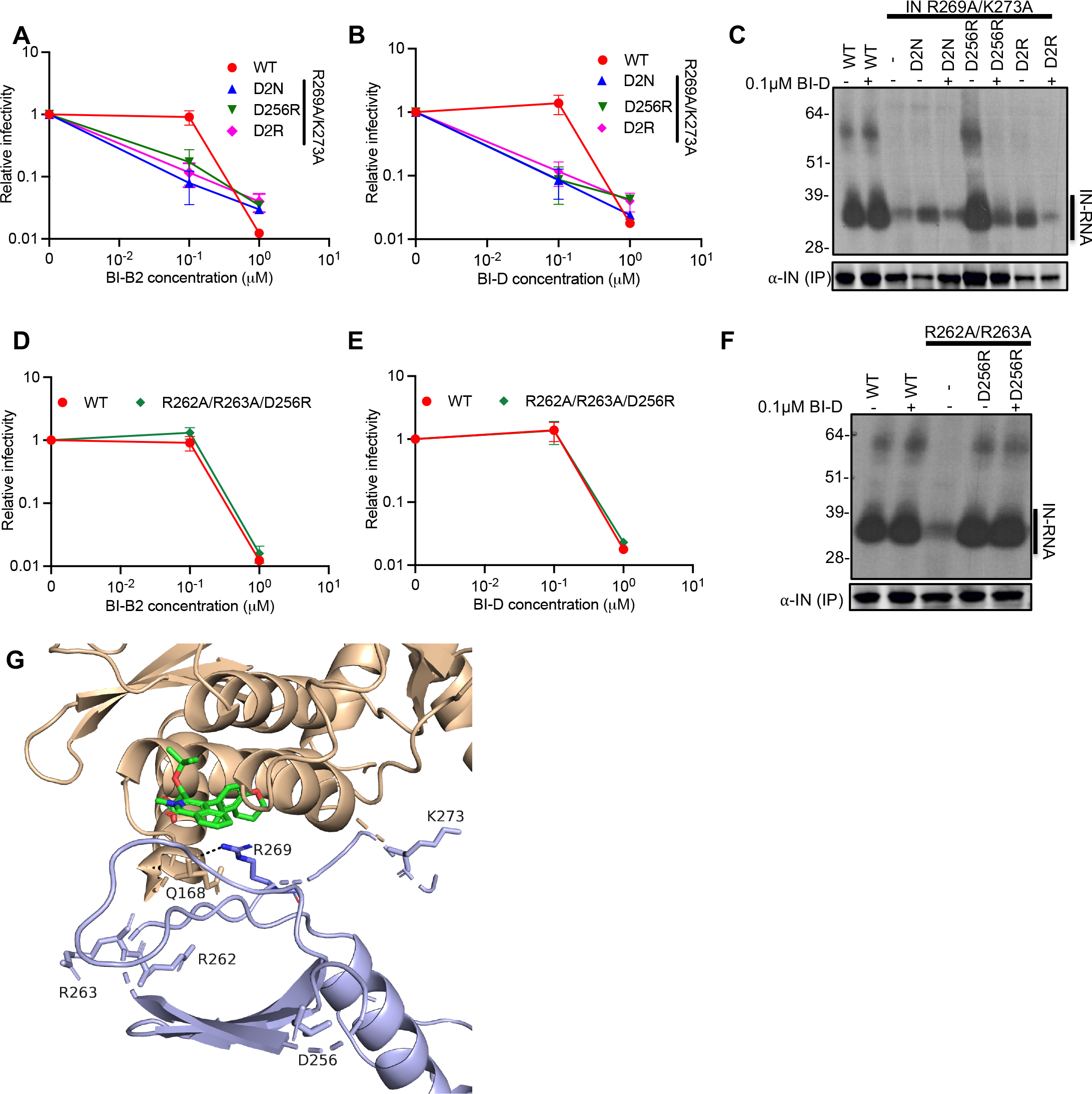
Secondary site suppressors of the IN R269A/K273A mutant increase susceptibility to ALLINIs. **(A, B, D, E)** Titers of viruses bearing the indicated substitutions in IN produced from HEK293T cells at different concentrations of ALLINIs, BI-B2 or BI-D. WT or IN mutant HIV-1_NL4-3_ in cell culture supernatants were titered on TZM-bl indicator cells. The titer values are represented relative to the mock control of each mutant (set to 1). Data are from two independent biological replicates. **(C, F)** Autoradiogram of IN-RNA adducts immunoprecipitated from WT or IN mutant HIV-1_NL4-3_ virions produced from HEK293T cells in the presence of 0.1μM of BI-D. The amount of immunoprecipitated IN protein was visualized by the immunoblot shown below. Immunoblots and CLIP autoradiographs results are a representative of three independent replicates. **(G)** The GSK-1264 binding pocket between CCD-CCD (brown) and CTD (light blue) from different IN subunits is shown (62) (PDB ID: 5HOT). GSK-1264 is in green with nitrogen and oxygen atoms colored blue and red, respectively. The side chain of R269 is shown with nitrogen atoms colored in dark blue. Interactions between CTD R269 and CCD Q168 are indicated by dashed lines.

### Characterization of IN mutations present in latently infected cells

Persistence of HIV-1 in memory CD4+ T-cells as latent proviruses constitutes a major barrier to HIV-1 cure. Although the majority of HIV-1 proviruses in these cells are defective (77), recent evidence suggests that defective proviruses can be transcribed into RNAs that are spliced, translated and can be recognized by HIV-1-specific cytotoxic T lymphocytes (78). We decided to characterize IN mutations isolated from latently infected cells, given the possibility that class II IN mutations existing in latently infected cells can result in the formation of defective particles that may subsequently modulate immune responses. Though relatively uncommon, we found the presence of R224Q, S230N, E246K and G272R substitutions in IN-CTD (**Fig. 9A**). Of note, only the R224Q substitution resulted in loss of a positive charge, whereas the E246K and G273R substitutions resulted in gain of positive charges. These mutations were introduced into the NL4-3 proviral backbone with minimal effects on Gag expression and particle release (**Fig. 9B**). Although the E246K virus was significantly less infectious (**Fig. 9C**), we did not find any evidence for loss of IN-gRNA binding (**Fig. 9D**), suggesting that this mutant likely displays a class I phenotype. Thus, we conclude that the class II mutant viruses are rarely present in the latently infected cells and therefore unlikely to contribute to chronic immune activation.

**Figure 9.**
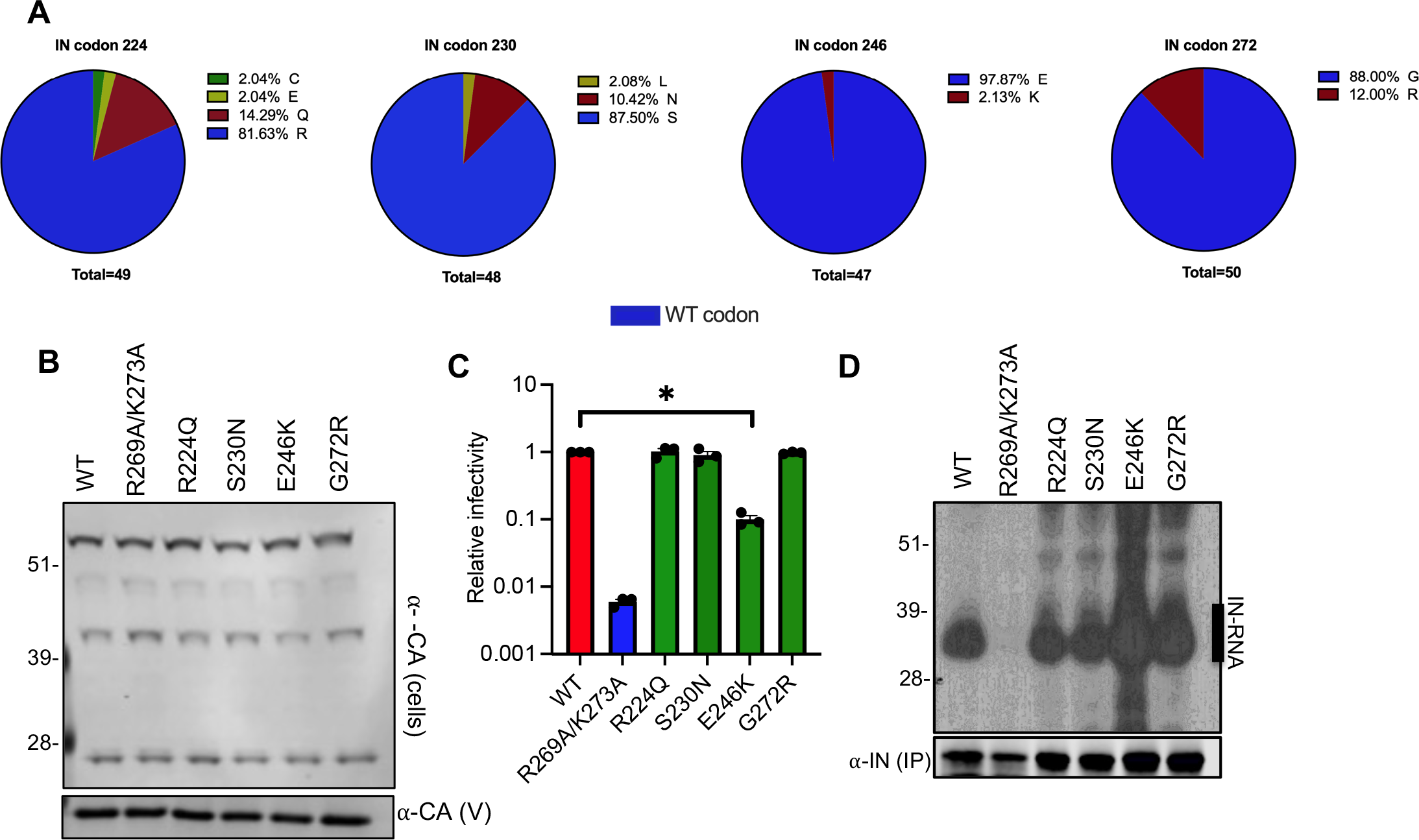
Characterization of IN mutations present in latently infected CD4+ T-cells. **(A)** Frequency of IN substitutions at amino acids 224, 230, 246 and 272 derived from HIV-1 sequences from latently infected CD4+ T-cells is shown. **(B)** HEK293T cells were transfected with proviral HIV-1_NL4-3_ expression plasmids carrying the R224Q, S230N, E246K, and G272R IN mutations. Cell lysates and virions were purified two days post transfection and analyzed by immunoblotting for CA and IN. The image is representative of two independent experiments. **(C)** WT or IN mutant HIV-1L4-3 viruses in cell culture supernatants were titered on TZM-bl indicator cells. The titers are presented relative to WT (set to 1). The columns represent the average of three independent experiments and the error bars represent SEM (*p<0.05, by one-way ANOVA multiple comparison test with Dunnett’s correction). **(D)** Autoradiogram of IN-RNA adducts immunoprecipitated from virions bearing the indicated substitutions in IN. Immunoblots below show the amount of immunoprecipitated IN.

## DISCUSSION

Class II IN mutations impair virion particle maturation by blocking IN-gRNA binding in virions and those within the CTD, including R269A/K273A and R262A/R263A substitutions, impede IN-gRNA binding without affecting functional oligomerization of IN (15, 38). During serial passaging of the IN R269A/K273A class II mutant virus, the D256N and D270N substitutions emerged sequentially. We also wondered whether mutations outside of IN, such as CA and NC, could also arise, given that the IN R269A/K273A mutant is still catalytically active and compensatory mutations in CA and NC could potentially provide alternative mechanisms for the proper packaging of the gRNA within virions. However, we did not observe the emergence of such substitutions showcasing the distinct role of IN-gRNA binding in proper virion maturation. Furthermore, the emergence of compensatory substitutions only in CTD, but not NTD or CCD of IN, highlights the importance of the IN-CTD in mediating direct RNA binding.

It is worth noting that D256N and D270N substitutions each arose through a single mutation (D256N: GAC AAC, D270N: GAU AAU), and thus likely provided an easier pathway for suppression than reverting back to R269 and K273, each of which would require two mutations. Likewise, D-to-R and D-to-K substitutions require two and three mutations, respectively, and thus likely had a higher barrier to arise in culture. Though other mutations in IN could in principle restore the ability of IN to bind RNA, rise of such mutations was likely additionally constrained in part by the necessity to maintain a catalytically active IN.

The IN-CTD is decorated by numerous acidic and basic residues (**Fig. 3A**). While the IN R269A/K273A substitutions resulted in the loss of 2 positive charges, the compensatory D2N substitutions restored the overall local charge within the CTD, suggesting that electrostatic interactions contribute to IN-CTD binding to the gRNA. Consistent with this hypothesis, the D256R, D278R and D279R substitutions that resulted in the gain of two positive charges locally were all sufficient to substantially increase or restore IN-gRNA binding for the IN R269A/K273A mutant. Furthermore, these observations extended to another class II mutant, IN R262A/R263A, whereby the D256R and to a lesser extent the D256K substitutions restored RNA binding and subsequent infectivity. In contrast, the D2N substitutions did not enhance RNA-binding or infectivity for the IN R262A/R263A mutant, demonstrating a degree of context/localization dependency. In other words, it is possible that the proximity of the D270N to R269A/K273A residues may explain why it was more effective in restoring RNA binding particularly for this class II mutant but not for IN R262A/R263A. Cumulatively, these findings strongly suggest an electrostatic component of IN-gRNA interactions. Inspection of the available X-ray structure (62) indicates that all IN residues implicated in RNA binding by the present study are positioned either in the highly flexible C-terminal tail (aa 261-275) or the 252-257 loop (**Fig. 5H**). The pliable nature of these CTD regions could be essential for allowing IN to optimally bind to cognate gRNA.

Arg and Lys residues are commonly employed by numerous RNA-binding proteins, and Arg residues are generally more heavily involved in interactions with all bases (70, 79–83). Though the electrostatic interactions between the Arg/Lys residues of IN-CTD and gRNA imply a level of non-specificity, IN binds to distinct locations on the gRNA in virions and displays high binding affinity to structured elements, such as TAR (38). Thus, it is likely that IN-gRNA interactions are mediated by both the non-specific interactions of the basic residues with the RNA phosphate backbone and specific interactions with the cognate RNA. For example, recognition of the TAR loop by Tat and the super elongation complex is based on a complex set of interactions: the super elongation complex primarily reads out the structure as opposed to the sequence, whereas the Tat arginine rich motif (ARM) interacts with the TAR bulge and the major groove through electrostatic interactions with the RNA backbone and H-bonding with specific bases (84, 85). Importantly, we note that while the present study elucidates crucial roles of the IN-CTD basic residues, other yet to be identified IN amino acids are likely to also contribute to high affinity binding of select gRNA segments. Structural studies of IN in complex with cognate RNA oligonucleotides are needed to tease apart the specificity determinants for IN-gRNA interactions.

RNA binding proteins commonly encode modular RNA-binding domains (i.e. RRMs and KH domains), which form specific contacts with short degenerate sequences (86–88). Utilization of multiple RRMs/KH domains is thought to create a much larger binding interface, which in turn allows for specific, high affinity binding to target RNAs (86, 87, 89). Functionally important IN tetramerization (15) may similarly enhance binding affinity and specificity to cognate gRNA elements. In this respect, it is worth noting that CTDs of the two inner protomers in the tetrameric intasome structure contact extensively with DNA, whereas the outer CTDs are positioned in close proximity to the inner CTDs and only partially contribute DNA binding (90). It remains to be seen whether the inner CTDs could also preferentially engage gRNA or CTD- nucleic acid interactions within IN-gDNA and IN-gRNA complexes differ substantially.

ALLINIs selectively bind at the IN CCD dimer interface at the LEDGF/p75 binding pocket and induce aberrant IN multimerization (12, 38), which is thought to underlie the mechanism by which they inhibit IN-gRNA binding. Prior biochemical and structural studies suggested that ALLINIs also engage the CTD of IN. For example, the quinoline-based ALLINI, GSK-1264, is buried between the CTD of one IN dimer and the CCD of a nearby dimer resulting in the formation of open, inactive IN polymers (62). Furthermore, mutation of Y226, W235, K264 and K266 residues within the CTD prevents ALLINI-induced aberrant multimerization of IN (62, 75, 76). However, evidence for CTD involvement in relevant infection settings is lacking, given that the mutation of the residues that mediate ALLINI induced hyper-multimerization of IN affect the catalytic activity as well as RNA binding properties of IN (62, 91, 92). Our findings fill this critical gap and provide key genetic evidence that ALLINI mechanism of action involves distinct residues within the CTD, namely R269 and K273. The X-ray structure of GSK-1264 induced IN polymers (62) reveals that R262, R263, K273A as well as D256 are distanced from the inhibitor binding pocket, whereas the side chain of R269 points toward the inhibitor bound at the CTD- CCD dimer interface (**Fig. 7G**). The guanidine group of R269 engages with Q168 of the CCD from another IN subunit and is positioned within 3 Å of GSK-1264. Yet, there is no interaction between R269 and the inhibitor as the positively charged guanidine group faces a hydrophobic part of the benzodihydropyran moiety of GSK-1264 (**Fig. 7G**). Conversely, the R269A substitution could provide more favorable hydrophobic environment for the CTD-ALLINI-CCD interactions that may explain its increased sensitivity to ALLINIs.

Overall, our studies reveal that pliable electrostatic interactions play an important role in mediating IN-CTD-gRNA interactions and demonstrate that CTD is a key determinant of both RNA binding and ALLINI sensitivity. IN-gRNA binding is essential for HIV-1 virion morphological maturation and infectivity, thus an excellent target for novel antiretroviral compounds. ALLINI- mediated inhibition of the non-catalytic function of IN can complement existing INSTI-based therapies and increase the barrier to drug resistance substantially. Of note, a highly potent and safe pyrrolopyridine-based ALLINI, STP0404, has recently advanced to human trials (65). Further structural characterization of IN-gRNA and IN-ALLINI complexes will be crucial to determine the precise rules that govern IN-gRNA interactions and guide the development of novel therapeutics that target IN-gRNA interactions.

## MATERIALS AND METHODS

### Plasmids and Compounds

IN mutations were introduced into the HIV-1_NL4-3_ full-length proviral plasmid (pNL4-3) by overlap extension PCR. Forward and reverse primers containing IN mutations were used in PCR reactions with forward (with EcoRI restriction endonuclease site) and reverse (with AgeI restriction endonuclease site) outer primers. The resulting fragments containing the desired mutations were mixed at a 1:1 ratio and overlapped subsequently using the outer primer pairs. The overlap fragments were digested with AgeI and EcoRI before cloning into pNL4-3 plasmids. The pLR2P-VprIN plasmid expressing a Vpr-IN fusion protein has been previously described (74). IN mutations were introduced in the pLR2P-VprIN plasmid using the QuikChange Site-Directed Mutagenesis kit (Agilent Technologies). The presence of the desired mutations and the absence of unwanted secondary changes was assessed by Sanger sequencing.

ALLINIs, BI-D and BI-B2, were synthesized by Aris Pharmaceuticals using previously published protocols (93, 94). Compounds were dissolved in DMSO at a final concentration of 10 mM and stored in -80C.

### Cell lines and viruses

Hela-derived TZM-bL cells that express CD4/CXCR4/CCR5 receptor/coreceptors and bear an HIV-1 LTR driven β-galactosidase reporter were obtained from the NIH AIDS Reagent Program. HEK293T cells (ATCC CRL-11268) and TZM-bL cells were cultured in Dulbecco’s modified Eagle’s medium supplemented with 10% fetal bovine serum. Human CD4+ T-cell line, MT4 (NIH AIDS Reagents), were cultured in RPMI 1640 medium supplemented with 10% fetal bovine serum. A derivative of MT4-LTR-GFP (MT4-GFP) indicator cells bearing an HIV-1 LTR driven GFP reporter have been described before (95) and similarly maintained in RPMI 1640 medium supplemented with 10% fetal bovine serum. All cell lines were obtained from the American Type Culture Collection and NIH AIDS Reagents where short tandem repeat (STR) profiling was performed. MT-4 T-cells were further authenticated by STR profiling at Washington University School of Medicine Genome Engineering and iPSC center. The cell lines are regularly inspected for mycoplasma contamination using MycoAlert mycoplasma detection kit (Lonza) and checked for being free of any other contaminations.

For generation of HIV-1 stocks, HEK293T cells grown in 10-cm dishes were transfected with the pNL4-3 and a vesicular stomatitis virus glycoprotein (VSV-G) expression plasmids at a ratio of 4:1 using polyethyleneimine (PolySciences, Warrington, PA). Cell culture supernatants containing infectious virions were collected two days post-transfection, filtered, aliquoted and stored at -80°C. For characterization of the effects of compensatory mutations on virus replication, HEK293T cells were transfected in 24-well plates with pNL4-3-derived plasmids similarly but without VSV-G pseudotyping. Cell lysates collected at 2 days post-transfection were analyzed by immunoblotting as detailed below. In parallel, cell culture supernatants containing virions were titered on TZM-bL cells using a β-galactosidase assay (Galactostar, Thermo Fisher) per manufacturer’s instructions. Virus replication in TZM-bL cells was limited to a single-cycle via addition of dextran sulfate (50 μg/mL) 6-14 hours post-infection. An aliquot of the viruses were also subjected to a qPCR-based reverse-transcriptase (RT) activity assay (96). Virus titers obtained from infection of TZM-bL cells was normalized to particle numbers based on values obtained from RT activity assays to determine the infectiousness (i.e. infectious units/particle) of viruses. A fraction of the cell culture supernatant collected from transfected HEK293T cells was also concentrated by centrifugation (16,100 x g, 4°C, 90min) on a 20% sucrose cushion prepared in 1x SDS-PAGE sample buffer for analysis of virion-associated proteins by immunoblotting as described below.

### Analysis of compensatory mutations

Compensatory substitutions that arose in the background of HIV-1_NL4-3 IN (R269A/K273A)_ class II IN mutant virus were isolated by serial passaging as follows. In brief, one million MT4-GFP cells were infected at a multiplicity of infection (MOI) 2 for the WT virus. An equivalent particle number (as determined by a qPCR-based reverse transcriptase (RT) activity assay (96)) of the HIV-1_NL4-3 IN (R269A/K273A)_ virus was used as inoculum. Infected cells were split at a ratio of 1:5 every 3-4 days and infections were monitored by microscopy as well as flow cytometry of GFP positive cells at the indicated intervals shown in Figure 1A. Cell culture supernatants containing virions were also collected at the same time intervals for subsequent sequencing analysis. At the end of each passage (i.e. when infections plateaued at >75%GFP+ cells), cell culture supernatants containing infectious virions were collected. The resultant viruses were first titered on MT4-GFP cells and used to infect MT4-GFP cells in the next passage at an MOI of 2 i.u./cell for WT and an equivalent particle number for the class II IN mutant virus (or its derivative bearing compensatory mutations). Aliquots of infected cells and viral particles in the cell culture supernatants were collected over the duration of each passage as above.

For sequencing analysis of viruses bearing compensatory mutations, virions collected over the three passages were concentrated on 20% sucrose cushions (prepared in 1X phosphate buffered saline (PBS)) by ultracentrifugation (28000 rpm, 4°C, 90min, Beckman SW41 rotor). Genomic RNA was isolated from pelleted virions using Trizol per manufacturer’s instructions. Extracted RNA was prepared for deep sequencing using the Illumina® TruSeq® Stranded Total RNA library kit following the manufacturer’s instructions but omitting the rRNA depletion step. Resulting libraries were sequenced on an Illumina HiSeq 2000 platform at the Genome Technology Access Center at Washington University School of Medicine. Sequencing reads were mapped to the pNL4-3 reference viral genome allowing for 2 mismatches using the Bowtie algorithm (i.e. -v 2, -m 10 parameters)(97) and the frequency of compensatory mutations was determined thereafter.

### Immunoblotting

Viral and cell lysates were resuspended in sodium dodecyl sulfate (SDS) sample buffer, separated by electrophoresis on Bolt 4-12% Bis-Tris Plus gels (Life Technologies) and transferred to nitrocellulose membranes (Hybond ECL, Amersham). The membranes were then probed overnight at 4°C with a mouse monoclonal anti-HIV p24 antibody (183-H12-5C, NIH AIDS reagents) or a mouse monoclonal anti-HIV integrase antibody (98) in Odyssey Blocking Buffer (LI-COR). Membranes were probed with fluorophore-conjugated secondary antibodies (LI-COR) and scanned using a LI-COR Odyssey system. IN and CA levels in virions were quantified using Image Studio software (LI-COR).

### Vpr-IN trans-complementation experiments

HIV-1_NL4-3_ bearing the class I IN mutation D116N was trans-complemented with class II mutant proteins as previously described (74). In brief, HEK293T cells plated in 24-well dishes were co-transfected with full-length proviral plasmids expressing HIV-1_NL4-3 IN (D116N)_, VSV-G, and derivatives of the pLR2P-VprIN plasmids bearing class II IN mutations (or the compensatory mutations thereof) at a ratio of 6:1:3 using polyethyleneimine as described above. Cell-free virions were collected from cell culture two days post-transfection. MT-4 T-cells were infected by the resultant virus stocks and the integration capability of trans-complemented class II IN mutants was measured by the yield of progeny virions in cell culture supernatants over a 6-day period using the aforementioned reverse transcriptase (RT) activity assay as described before (74). Briefly, MT-4 T-cells were incubated with virus inoculum in 96 V-bottom well plates for 4hr at 37°C before washing away the inoculum and replacing it with fresh media. Right after the addition of fresh media and over the ensuing 6 days, an aliquot of the media was collected and the amount of virions in culture supernatant was quantified by measuring a qPCR-based RT activity using assay (96), as also described above.

### CLIP experiments

CLIP experiments were conducted as previously described (15, 72, 73, 99, 100). In short, cells in 15 cm cell culture plates were transfected with 30 μg full-length proviral plasmid (pNL4-3) DNA or derivatives carrying the IN mutations. 4-thiouridine (4SU) was added to the cell culture media for 16hr before virus harvest. Cell culture supernatants containing virions were filtered through 0.22 μm filters and pelleted by ultracentrifugation through a 20% sucrose cushion prepared in 1X phosphate-buffered saline (PBS) using a Beckman SW32-Ti rotor at 28,000rpm for 1.5hr at 4°C. The virus pellets were resuspended in 1XPBS and UV-crosslinked for two consecutive times at an energy setting of 500 mJ in a Boekel UV-crosslinking chamber equipped with UV368 nm bulbs. Following lysis in 1X RIPA buffer, IN-RNA complexes were immunoprecipitated using a mouse monoclonal anti-IN antibody (98). Bound RNA was end-labeled with -^32^

P-A TP and T4 polynucleotide kinase. The isolated protein-RNA complexes were separated by SDS-PAGE, transferred to nitrocellulose membranes (Hybond ECL, Amersham), and exposed to autoradiography films to visualize IN-RNA complexes. If needed, following immunoblotting and quantitation of immunoprecipitated IN, equivalent amounts of immunoprecipitated IN were re-run to accurately assess the ability of IN variants to bind RNA. Autoradiographs were quantitated by the ImageStudio software (LI-COR). Lysates and immunoprecipitates were also analyzed by immunoblotting using the aforementioned mouse monoclonal antibodies against IN.

### IN multimerization in virions

HEK293T cells grown in 10-cm dishes were transfected with 10 μg pNL4-3 plasmid DNA bearing WT IN or the indicated *pol* mutations within IN coding sequence as above. Two days post-transfection, cell-free virions in cell culture supernatants were pelleted through a 20% sucrose cushion prepared in 1xPBS using a Beckman SW41-Ti rotor at 28,000 rpm for 1.5hr at 4°C. Pelleted virions were resuspended in 1xPBS and treated with a membrane-permeable crosslinker, EGS (ThermoFisher Scientific), at a concentration of 1mM for 30 min at room temperature. Crosslinking was stopped by the addition of SDS-PAGE sample buffer. The cross- linked samples were then separated on 3-8% Tris-acetate gels and analyzed by immunoblotting using a mouse monoclonal anti-IN antibody (98).

### Virus production and transmission electron microscopy

HEK293T cells grown in 15-cm plates were transfected with 30 μg full-length proviral plasmid (pNL4-3) DNA containing WT IN or the indicated *pol* mutations within IN coding sequence as above. Two days post transfection, cell culture supernatants were filtered through 0.22 μm filters, and pelleted by ultracentrifugation using a Beckman SW32-Ti rotor at 28,000 rpm for 1.5 hr at 4°C. Fixative (2% paraformaldehyde/2.5% glutaraldehyde (Polysciences Inc., Warrington, PA) in 100 mM sodium cacodylate buffer, pH 7.2) was gently added to resulting pellets, and samples were incubated overnight at 4°C. Samples were washed in sodium cacodylate buffer and postfixed in 1% osmium tetroxide (Polysciences Inc.) for 1 hr. Samples were then rinsed extensively in dH_2_0 prior to en bloc staining with 1% aqueous uranyl acetate (Ted Pella Inc., Redding, CA) for 1 hr. After several rinses in dH_2_0, samples were dehydrated in a graded series of ethanol and embedded in Eponate 12 resin (Ted Pella Inc.). Sections of 95 nm were cut with a Leica Ultracut UCT ultramicrotome (Leica Microsystems Inc., Bannockburn, IL), stained with uranyl acetate and lead citrate, and viewed on a JEOL 1200 EX transmission electron microscope (JEOL USA Inc., Peabody, MA) equipped with an AMT 8 megapixel digital camera and AMT Image Capture Engine V602 software (Advanced Microscopy Techniques, Woburn, MA). 100 virions from each sample was classified for virion morphology.

### Size exclusion chromatography (SEC)

All of the indicated mutations were introduced into a plasmid backbone expressing His_6_ tagged pNL4-3-derived IN by QuikChange site directed mutagenesis kit (Agilent) (66). His_6_ tagged recombinant pNL4-3 WT and mutant IN molecules were expressed in BL21 (DE3) *E. coli* cells followed by nickel and heparin column purification as described previously (66, 101). Recombinant WT and mutant IN molecules were analyzed on Superdex 200 10/300 GL column (GE Healthcare) with running buffer containing 20 mM HEPES (pH 7.5), 1 M NaCl, 10% glycerol and 5 mM BME at 0.3 mL/min flow rate. The proteins were diluted to 10 µM with the running buffer and incubated for 1 h at 4°C followed by centrifugation at 10,000xg for 10 min. Multimeric form determination was based on the standards including bovine thyroglobulin (670,000 Da), bovine gamma-globulin (158,000 Da), chicken ovalbumin (44,000 Da), horse myoglobin (17,000 Da) and vitamin B12 (1,350 Da). Retention volumes for different oligomeric forms of IN were as follows: tetramer ∼12.5 mL, dimer ∼14 mL, monomer ∼15-16 mL.

### Analysis of IN-RNA binding *in vitro*

To monitor IN-RNA interactions we utilized AlphaScreen-based assay (38), which allows to monitor the ability of IN to bind and bridge between two TAR RNAs. Briefly, equal concentrations (1 nM) of two synthetic TAR RNA oligonucleotides labeled either with biotin or DIG were mixed and then streptavidin donor and anti-DIG acceptor beads at 0.02 mg/mL concentration were supplied in a buffer containing 100 mM NaCl, 1 mM MgCl_2_, 1 mM DTT, 1 mg/mL BSA, and 25 mM Tris (pH 7.4). After 2 hr incubation at 4°C, 320 nM IN was added to the reaction mixture and incubated further for 1.5 hr at 4°C. AlphaScreen signals were recorded with a PerkinElmer Life Sciences Enspire multimode plate reader.

### Structural modeling of IN

Electrostatic potential maps of WT and mutant IN CTDs were created by Adaptive Poisson-Boltzmann Solver (APBS) program (102) with macromolecular electrostatic calculations performed in PyMOL. The published crystal structure (62) (PDB ID: 5HOT) was used as a template. The calculation results are displayed as an electrostatic potential molecular surface. The low, mid, and high range values are -5, 0, and 5, respectively.

### Analysis of IN mutations from latently infected cells

Sequences identified from latently infected CD4+ T-cells (77) were downloaded from NCBI GenBank based on their accession numbers (KF526120-KF526339). Sequences were imported into the GLUE software framework (103, 104) and aligned. Multiple sequence alignments (MSAs) containing subtype B sequences were constructed using MUSCLE, manually inspected in AliView (105) and imported into a GLUE project database. Within GLUE, MSAs were constrained to the pNL4-3 reference to establish a standardized coordinate space for the gene being analyzed. Amino acid frequencies at each alignment position were summarized using GLUE’s amino-acid frequency calculation algorithm, which accounts for contingencies such as missing data and incomplete codons.

## ACKNOWLEDGEMENTS

This work was supported by NIH grants AI150497 (SBK), AI1508470 (U54 Center for HIV RNA Studies, SBK, RG), R01 AI143649 (MK) and Milton Schlesinger Student Fellowship (CSM). We thank all members of the Kutluay lab for critical suggestions and feedback and Wandy Beatty at Washington University for assistance and expertise in transmission electron microscopy.

## SUPPLEMENTAL FIGURE LEGENDS

Figure S1. Effect of compensatory D256N and D270N substitutions on viral gene expression, Gag processing, particle release and infectivity for the WT and HIV-1_NL4-3 IN (R269A/K273A)_ class II mutant virus. HEK293T cells were transfected with full-length proviral pNL4- 3 expression plasmids carrying mutations coding for the IN D256N, D270N and D256N/D270N (D2N) substitutions introduced on the WT IN and IN R269A/K273A backbones as described in Figure 1C. Cell lysates and purified virions were analyzed by immunoblotting for CA and IN. Quantification of unprocessed Gag-Pr55 (A), Gag processing intermediates (B-D), virion-associated IN levels (E) from 4 independent biological replicate experiments is shown. Error bars show the SEM. (F, G) Cell culture supernatants containing viruses were collected and used in an RT activity assay (F) or titered on TZM-bl indicator cells (G), whereby virus replication was limited to a single cycle by addition of dextran sulfate (50 μg/mL). The RT activity and titer values were normalized relative to WT (set to 1).

Figure S2. D256N and D270N substitutions restore RNA binding for the HIV-1_NL4-3 IN (R269A/K273A)_ class II mutant virus. IN-RNA adducts were immunoprecipitated from virions bearing the indicated substitutions in IN as in Fig. 2A. Autoradiography images were quantified for IN-RNA adducts and presented relative to WT (set to 1). Data are derived from three independent biological replicates with error bars showing the SEM. (****p<0.0001, ns=non-significant by one-way ANOVA multiple comparison test with Dunnett’s correction).

Figure S3. Effect of compensatory D256R and D270R substitutions on viral gene expression, Gag processing, particle release, infectivity and IN-RNA binding for the WT and HIV-1_NL4-3 IN (R269A/K273A)_ class II mutant viruses. HEK293T cells were transfected with full-length proviral pNL4-3 expression plasmids carrying mutations coding for the IN D2N, D256R, D270R and D256R/D270R (D2R) substitutions introduced on the IN R269A/K273A backbone as described in Fig. 3B, C. Cell lysates and purified virions were analyzed by immunoblotting for CA and IN. Quantification of unprocessed Gag-Pr55 (A), Gag processing intermediates (B-D), virion-associated IN levels (E) from 4 independent biological replicate experiments is shown. Error bars show the SEM. (F, G) Cell culture supernatants containing viruses were collected and used in an RT activity assay (F) or titered on TZM-bl indicator cells (G), whereby virus replication was limited to a single cycle by addition of dextran sulfate (50 μg/mL). The RT activity and titer values were normalized relative to WT (set to 1). (H) IN-RNA adducts were immunoprecipitated from virions bearing the indicated substitutions in IN as in Fig. 3D. Autoradiography images were quantified for IN-RNA adducts and presented relative to WT (set to 1). Data are derived from three independent biological replicates with error bars showing the SEM. (***p<0.001, **p<0.01 by one-way ANOVA multiple comparison test with Dunnett’s correction).

Figure S4. Effect of compensatory substitutions on viral gene expression, Gag processing, particle release, infectivity and IN-RNA binding for the WT and HIV-1_NL4-3 IN (R262A/R263A)_ class II mutant viruses. HEK293T cells were transfected with full-length proviral pNL4-3 expression plasmids carrying mutations coding for the IN D256N, D270N, D2N, D256K and D256R substitutions introduced on the WT IN or IN R262/R263A backbones as described in Fig. 4. Cell lysates and purified virions were analyzed by immunoblotting for CA and IN. Quantification of unprocessed Gag-Pr55 (A, G), Gag processing intermediates (B, C, D, H, I, J), virion-associated IN levels (E, K) from 4 independent biological replicates is shown. (F, L, M) Cell culture supernatants containing viruses were collected and titered on TZM-bl indicator cells (F, M) or used in an RT activity assay (L) to assess particle release. The RT activity and titer values were normalized relative to WT (set to 1). (N) IN-RNA adducts were immunoprecipitated from virions bearing the indicated substitutions in IN as in Fig. 4E. Autoradiography images were quantified and presented relative to WT (set to 1). Data are derived from three independent biological replicates with error bars showing the SEM. (****p<0.0001, **p<0.01, *p<0.05, ns=not significant by one-way ANOVA multiple comparison test with Dunnett’s correction).

