## Supplementary figures and images for "Emergence of compensatory mutations reveal the importance of electrostatic interactions between HIV-1 integrase and genomic RNA"

### Supplementary Figs 1-4

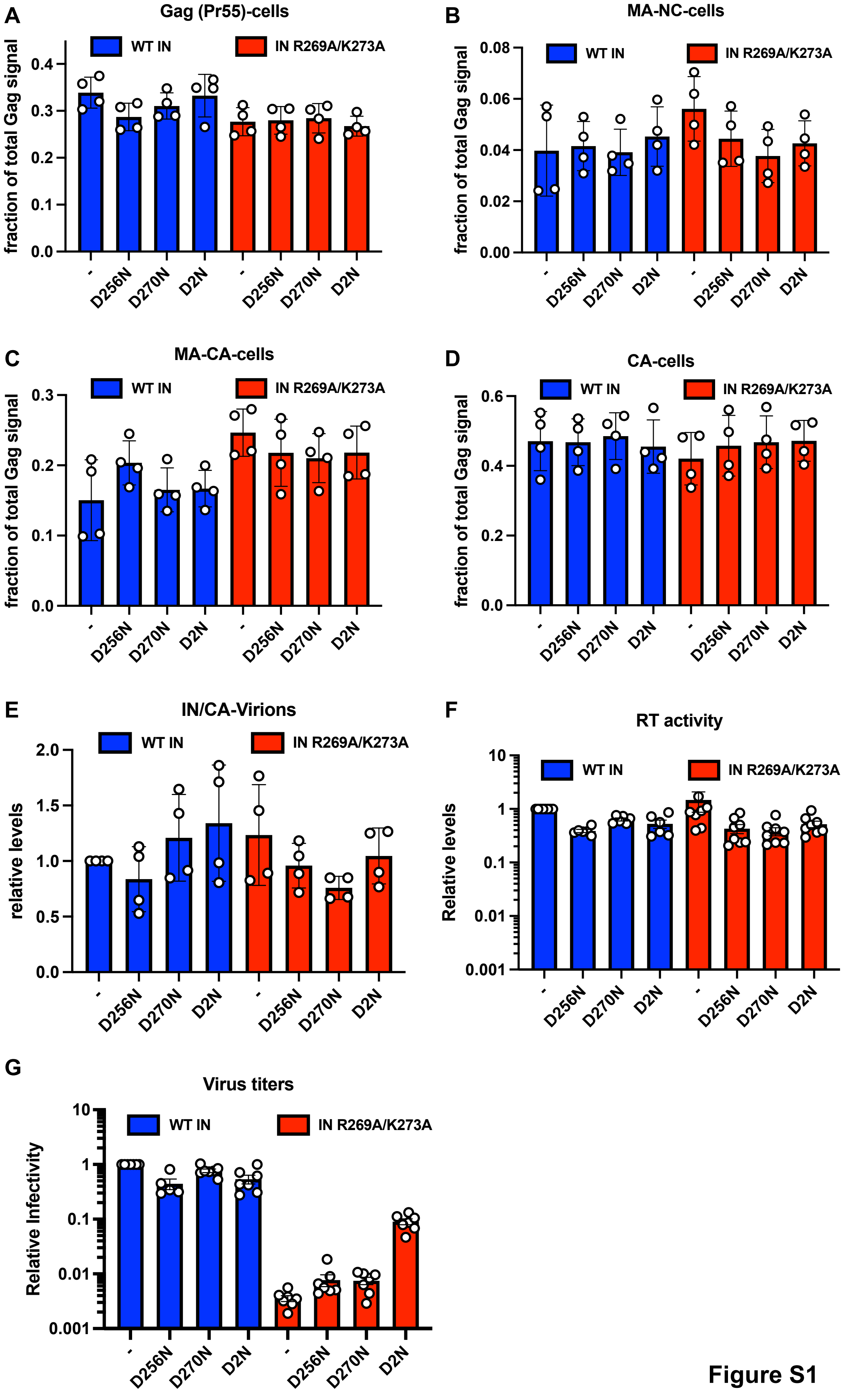

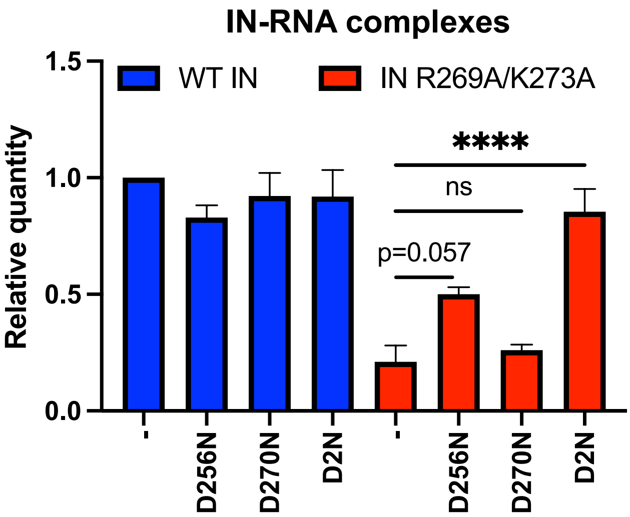

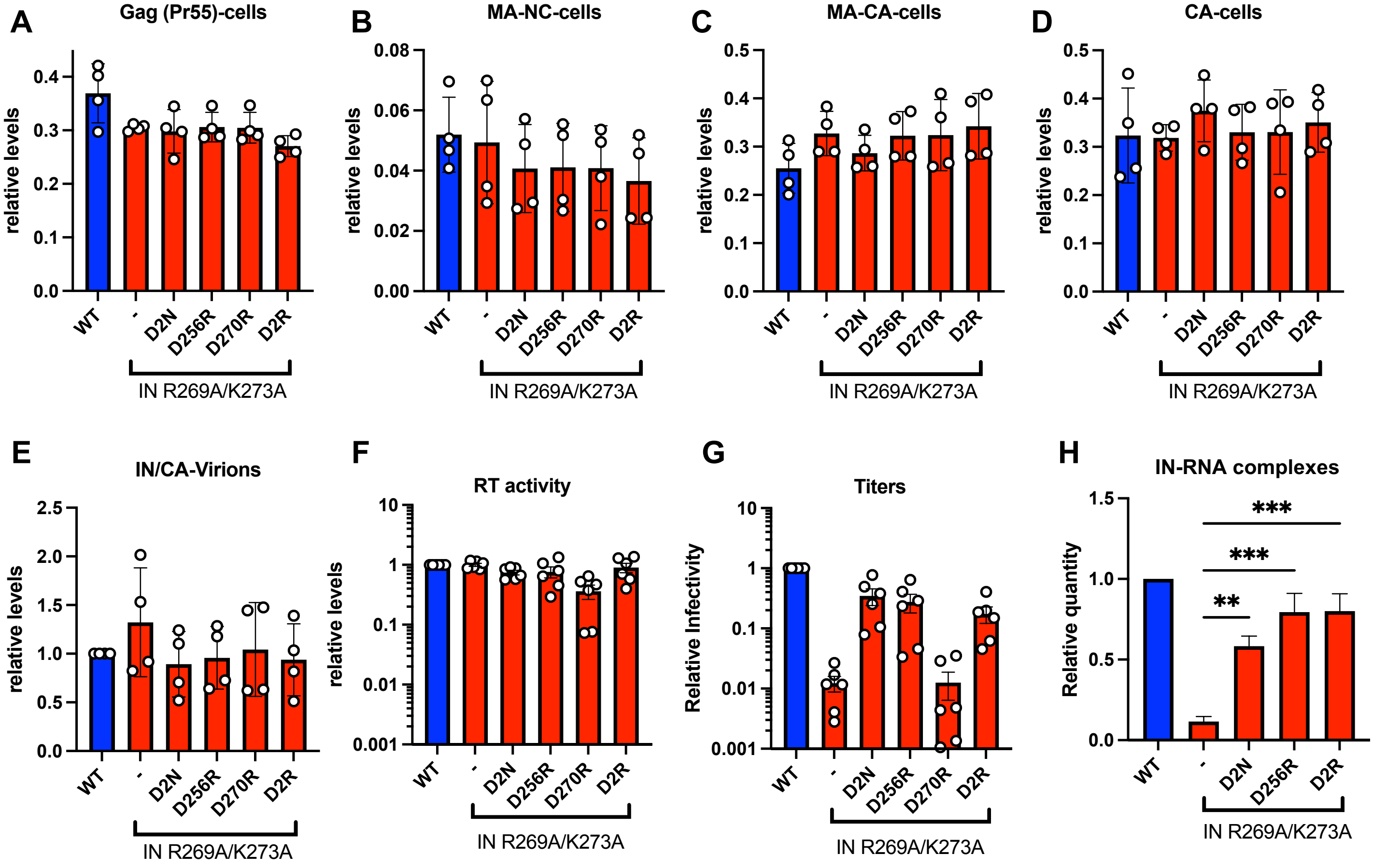

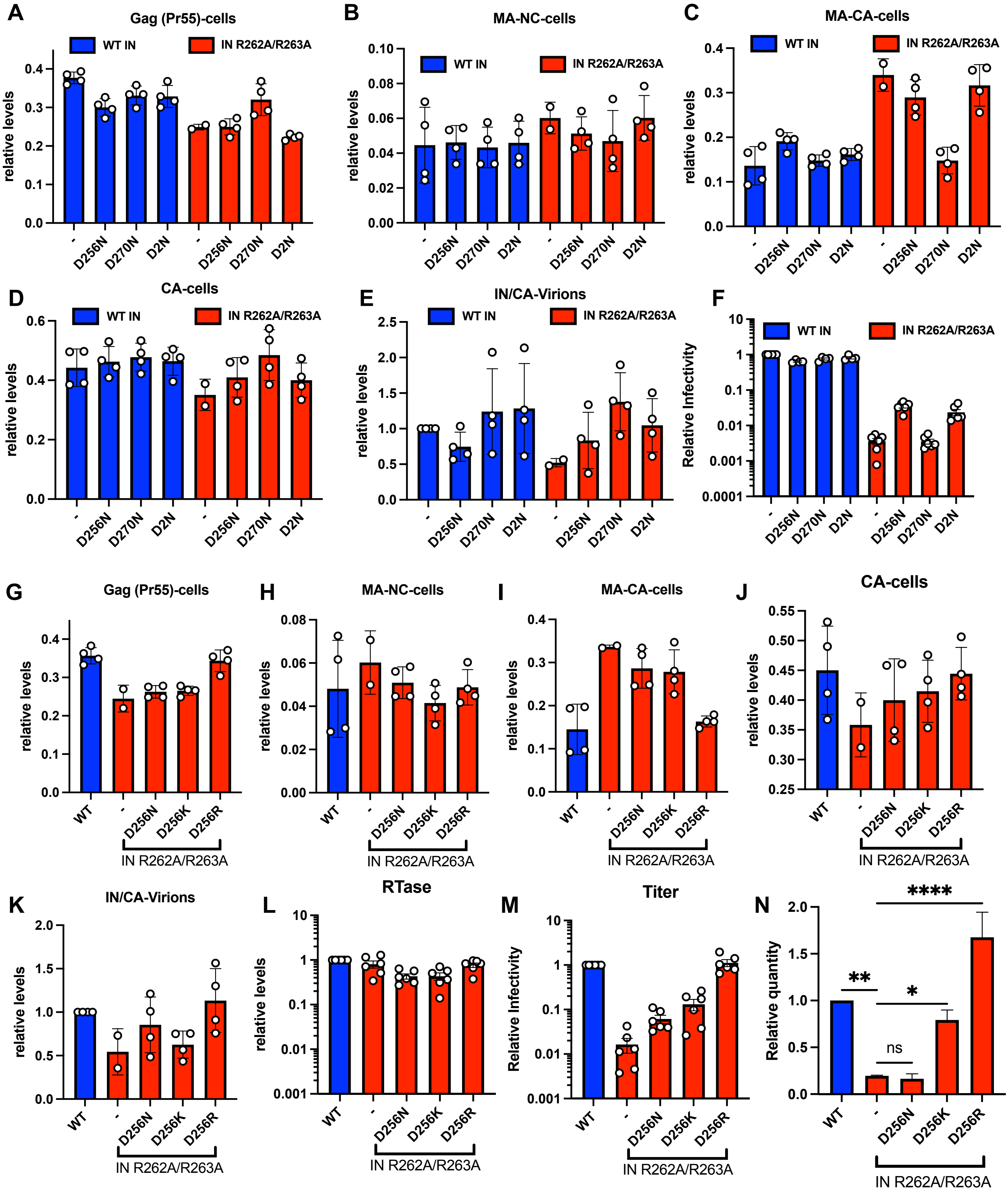
